# Machine Learning Assisted Spectral Fingerprinting for Immune Cell Phenotyping

**DOI:** 10.1101/2024.03.05.583608

**Authors:** Aceer Nadeem, Sarah Lyons, Aidan Kindopp, Amanda Jamieson, Daniel Roxbury

## Abstract

Spectral fingerprinting has emerged as a powerful tool, adept at identifying chemical compounds and deciphering complex interactions within cells and engineered nanomaterials. Using near-infrared (NIR) fluorescence spectral fingerprinting coupled with machine learning techniques, we uncover complex interactions between DNA-functionalized single-walled carbon nanotubes (DNA-SWCNTs) and live macrophage cells, enabling *in situ* phenotype discrimination. Through the use of Raman microscopy, we showcase statistically higher DNA-SWCNT uptake and a significantly lower defect ratio in M1 macrophages as compared to M2 and naïve phenotypes. NIR fluorescence data also indicate that distinctive intra-endosomal environments of these cell types give rise to significant differences in many optical features such as emission peak intensities, center wavelengths, and peak intensity ratios. Such features serve as distinctive markers for identifying different macrophage phenotypes. We further use a support vector machine (SVM) model trained on SWCNT fluorescence data to identify M1 and M2 macrophages, achieving an impressive accuracy of > 95%. Finally, we observe that the stability of DNA-SWCNT complexes, influenced by DNA sequence length, is a crucial consideration for applications such as cell phenotyping or mapping intra-endosomal microenvironments using AI techniques. Our findings suggest that shorter DNA-sequences like GT_6_ give rise to more improved model accuracy (> 87%) due to increased active interactions of SWCNTs with biomolecules in the endosomal microenvironment. Implications of this research extend to the development of nanomaterial-based platforms for cellular identification, holding promise for potential applications in real time monitoring of *in vivo* cellular differentiation.

**TOC Graphic:** 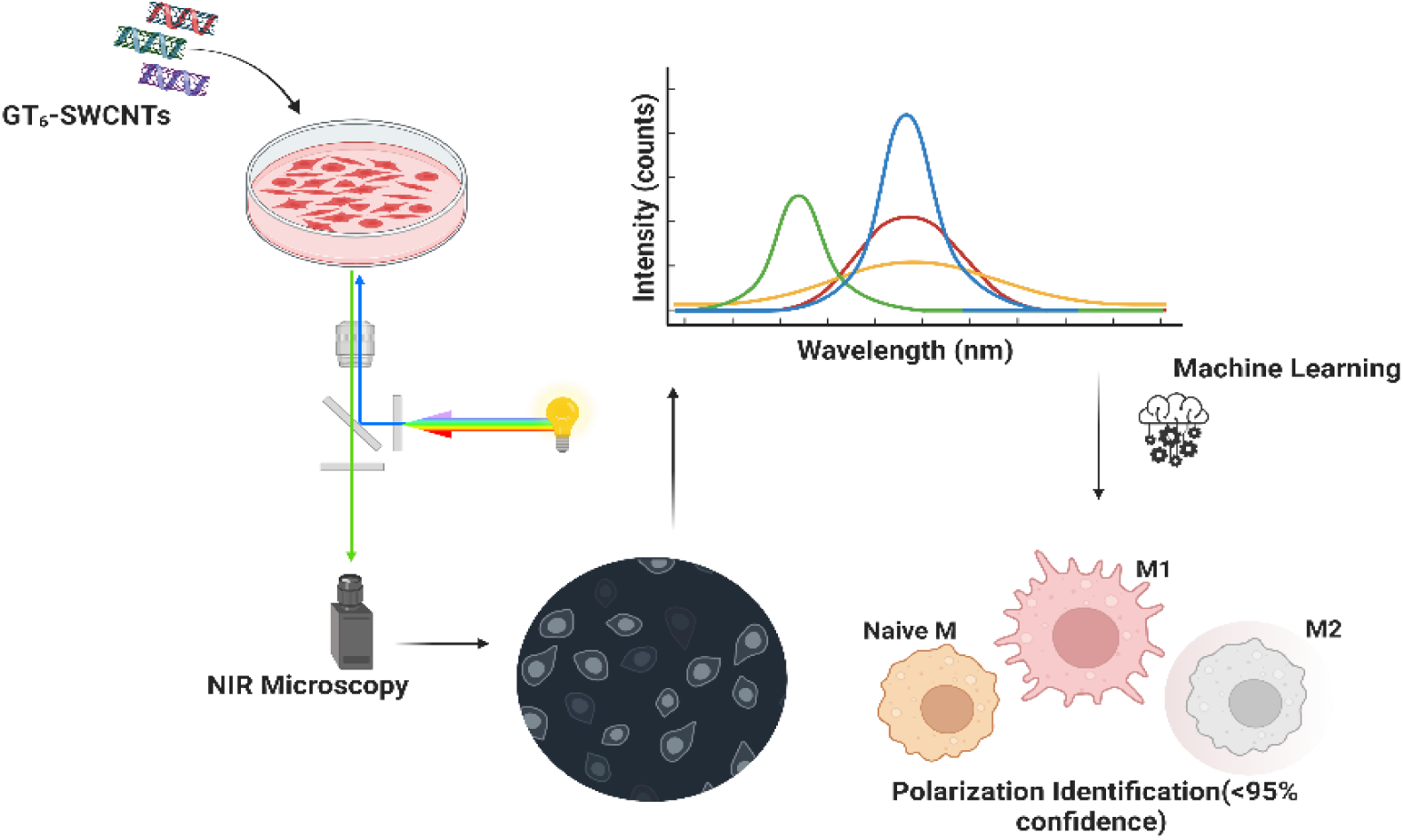

---

Biosensing is a rapidly emerging sector of the biomedical engineering field, specifically in human healthcare due to the non-invasive nature and tunable specificity of recent sensors. Typical examples of biosensors include immunosensors,^1^ DNA biosensors,^2^ enzyme-based biosensors,^3^ thermal and piezoelectric biosensors^4^ and optical biosensors,^5^ which utilize affinity towards antigens, chemical interactions, thermal fluctuations, affinity interactions and fluorescence, respectively, to pinpoint the presence or concentration change of a specific biological marker.^6,7^ Certain optical biosensors are of particular interest as they implement a light source to probe changes within cellular environments that can cause shifts in fluorescence wavelength, intensity, and/or spectral bandwidth which can be attributed to distinct cellular microenvironments.^8,9,10,11^

Single-walled carbon nanotubes (SWCNTs) have been successfully utilized as *in vitro* optical biosensors due to their intrinsic fluorescence and photostability.^12,13^ Moreover, the fluorescence of SWCNTs resides within the near-infrared (NIR) spectrum, a region which proves useful in biological imaging applications due to limited tissue absorbance, scattering, and autofluorescence.^14,15,16^ SWCNTs can be non-covalently functionalized with biocompatible polymers and biomolecules to increase environmental compatibility. Specifically, there has been success with different types of amphiphilic functionalization such as single-stranded DNA (ssDNA) and polyethylene glycol (PEG)-lipid conjugate wrappings.^17,18,19^ Through appropriate biocompatible functionalization, SWCNTs can be effectively internalized into mammalian cells via energy dependent endocytosis and phagocytosis, where they then enter the endosomal pathway and eventually localize within the lysosomes.^20,21^ The ssDNA wrapping of SWCNTs proves to be biocompatible as various studies have shown endocytosis and endosomal escape from different cell lines. We have previously shown that DNA sequence length plays a significant role in determining endocytosis and retention durations of DNA-functionalized single-walled carbon nanotubes (SWCNTs) within mammalian cells.^22^

Macrophages are immune cells of high interest in the field of biomedicine due to their implication in immune responses, inflammation regulation, and tissue repair. Macrophages play a crucial role in various biomedical contexts, such as host defense against pathogens, clearance of cellular debris, modulation of immune response and wound healing.^23^ Their versatility makes them a key player in understanding and addressing diseases, including infections, autoimmune disorders, and cancer. Macrophages are capable of polarizing and differentiating from monocytes and a naïve state into, broadly characterizing, either pro-inflammatory M1 phenotypes or pro-healing M2 phenotypes.^24,25^ These phenotypical changes are induced by signaling molecules and cytokines in the local cellular microenvironment.^26,27^ The inflammatory responses performed by M1 macrophages are dominated by toll-like receptor (TLR) and interferon signaling and can be polarized *in vitro* with interferon gamma (IFN-γ), tum or necrosis factor alpha (TNF-α), and lipopolysaccharide (LPS).^28,29^ In contrast, M2 macrophages are found in the proliferation and remodeling phases of wound healing, secreting cytokines to actively promote repair and recruit various cell types to clear cellular debris.^30,31^ Typical cytokines implicated in the differentiation of M2 macrophages are interleukin-4 (IL-4) and interleukin-10 (IL-10).^32,33^

The shift from inflammation to proliferation represents a pivotal stage in the wound healing process. An imbalance in the macrophage phenotype environment and the inability to transition between M1 and M2 states can lead to ulcers and chronic wounds.^34,35^ The wound healing process is similar to the human body’s reaction to diseases, such as cancers. In certain cancers, such as colorectal cancer, tumor cells will utilize macrophage anti-inflammatory characteristics to promote tumor growth, thus progressing the spread of cancer.^36^ Due to the vast differences between macrophage polarization states, there must be tight control of differentiation to avoid prolonged periods of inflammation. However, in the presence of chronic inflammation, imbalances of M1 and M2 phenotypes are observed and ultimately trigger changes in macrophage behavior.^37^ By quantifying and preventing changes in macrophage polarization states, there is potential for a decrease in disease progression.^38^

To identify M1 and M2 macrophage phenotypes and quantify any potential imbalance, in both wound healing and other affected disease, flow cytometry,^37^ surface marker analysis,^39^ and visual morphological confirmation via optical microscopy are frequently employed.^40,41^ Flow cytometry serves as a method which can rapidly differentiate different cells in a sample with high-throughput analysis. However, this method is not well adapted to identify different subsets of macrophage phenotypes as it requires a diverse array of surface markers and dyes which become prohibitively expensive.^42^ Moreover, issues in data collection can arise when samples have been exposed to fibrosis or other pathological changes, which is common in wound sites.^43^ The visual analysis of macrophage phenotypes via optical microscopy and immunofluorescence imaging can be useful in identifying M1 and M2 macrophages but becomes less reliable when comparting naïve macrophages to M2 due to their similar morphologies.^44^ Consequently, immunofluorescence microscopy has its own limitations.^45^

Morphological identification of samples through machine learning on cell shape and size, has been proven to accurately predict what subtype of M1 or M2 macrophage,^43^ however the identification of specific types of cells within a diseased and heterogenous sampling would be better served by a more robust method for classification. In a previous study, we have shown that DNA-SWCNTs can be used to map intracellular processes based on modulations in their Raman spectra.^46^ These differences induced throughout a single cell could be attributed to changes in the local environment of the SWCNTs, such as pH, salt concentration, and/or protein interactions. Here, utilizing NIR fluorescence data of SWCNTs within macrophages of varying phenotype coupled with machine learning, we show that a similar spectral fingerprinting method can be developed to accurately identify live M1, M2, and naïve macrophages. We find that DNA-SWCNTs are rapidly internalized into all macrophage phenotypes, with M1 demonstrating the highest rate of uptake. Once inside, we observe interesting trends in the Raman spectra of the SWCNTs as well as a DNA-sequence specific modulation in NIR fluorescence. We uncover that SWCNTs dispersed with a short DNA sequence (i.e. GT_6_-SWCNTs) are able to undergo the largest degree of SWCNT chirality-dependent modulations in NIR fluorescence, enabling the highest accuracy of *in vitro* discrimination between macrophage phenotypes. The presented SWCNT spectral fingerprinting coupled machine learning method is versatile and can be applied to other live-cell discrimination analyses.

## Results/Discussion

A model mammalian macrophage cell line (RAW 264.7 murine macrophages) was employed to investigate characteristic differences between Naïve (non-polarized, “NM”), M1, and M2 macrophage phenotypes. Cell type specific cytokines were added to NM cells to artificially stimulate them into differentially activated phenotypes, M1 and M2. Figure 1a represents transmitted light images of each macrophage phenotype captured through bright field microscopy. Upon closer examination, noticeable differences in size (figure S1) and morphology are evident (figure S2). M2 display a nearly round and circular morphology, like that of naïve macrophages, whereas M1 are larger in size, as well as lacking a well-defined shape. These distinct features offer a visual confirmation that we have different species of macrophages. Figure 1b shows a statistical bar graph highlighting the significant difference in cell area and size between the three cell species, these area and sizes were determined using image analysis as detailed in figure S3. Where M1 exhibit the largest average cell area measuring approximately 520 µm^2^, M2 and NM are observed to be considerably smaller in size with an average area of 200 µm^2^ (61% smaller) for M2 and 240 µm^2^ (56% smaller) for naïve macrophages.

**Figure 1.**
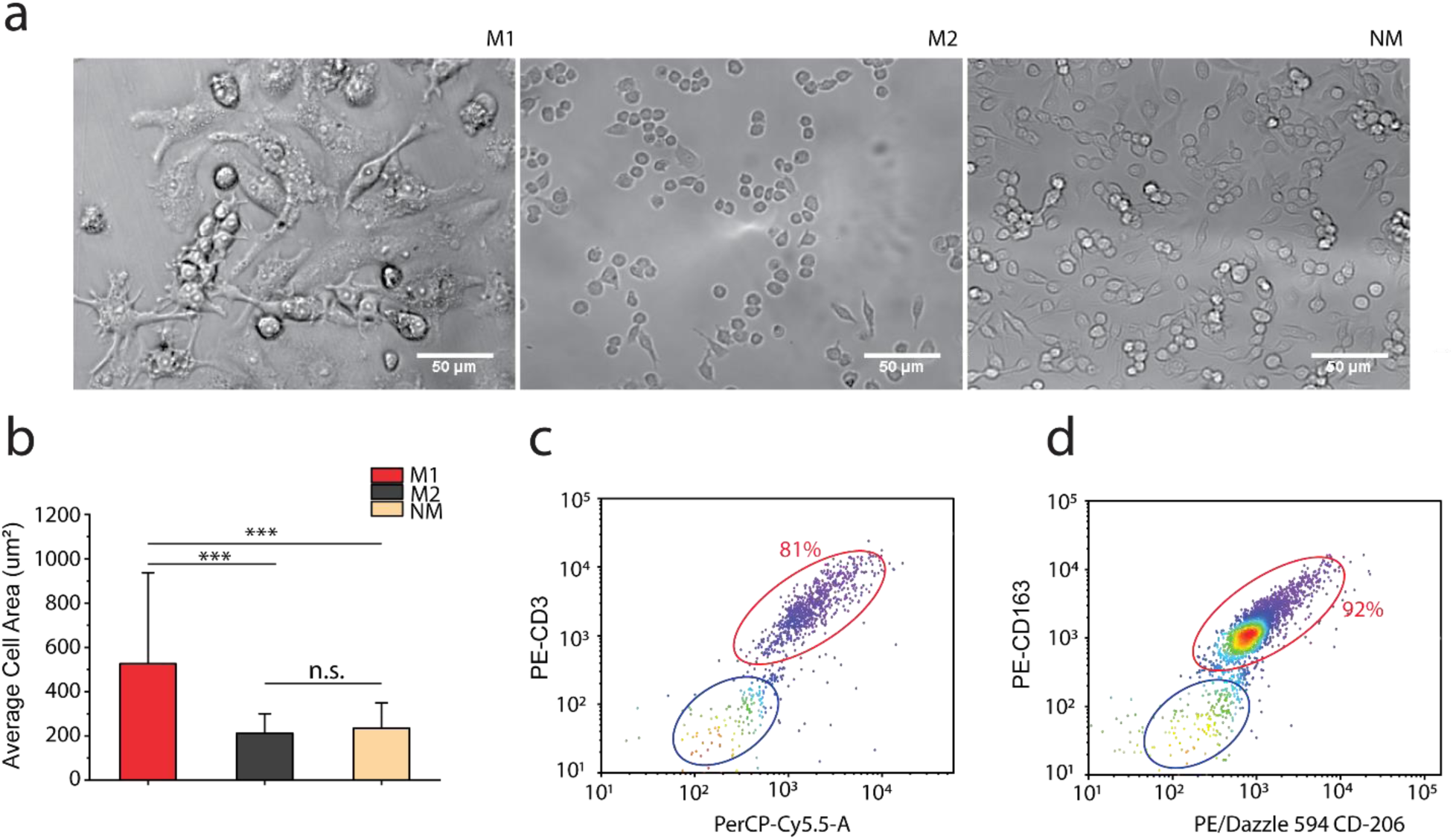
Cell Characterization. (a) Transmitted light images of M1, M2 and naïve macrophages, with scale bar of 50 µm. (b) Bar graph for average cell area, bars represent the average, and whiskers represent mean ± s.d. for each condition (*n* ≥ 500 cells per condition). Two-Sample t-test hypothesis testing analysis was performed between different samples (\*\*\**p* < 0.001). Fluorescent surface marker dependent flow cytometry density plot for (c) M1 type macrophages (d) M2 type macrophages, for each condition n ≥ 5×10^5^ cells/ml.

Following a 24-hour cytokine dose, fluorescence-activated cell sorting flow cytometry (FACS) was conducted to further validate the macrophage phenotypes. In figure 1c and 1d, fluorescent marker-specific cell density plots are presented, specifically gated to illustrate the population of M1 and M2 in the sample. Analysis of both scatter plots reveal that after the 24-hour cytokine dose, 81% of the cells in the sample exhibit M1 phenotype, while 92% display M2 macrophage phenotype. It is crucial to emphasize that the lower percentage of M1 population is primarily attributed to a substantial amount of cell death observed during the processing of M1 cells for FACS (see methods). This observed cell death can be directly linked to the distinct cell-surface adhesion characteristics of the two macrophage phenotypes. Classically activated M1 macrophages exhibit robust adhesion properties in contrast to the low-adherent nature of M2 macrophages.^47^ Consequently, the M1 sample shows increased cell debris and cell death due to the more adherent nature of M1 activated cells, this is further confirmed in an apoptosis/necrosis assay (figure S4). These results highlight the challenges associated with the differential adhesion properties and subsequent cell recovery during experiments for this study.

It is imperative to recognize and address the inherent error in the data moving forward. This margin of error stems from challenges in achieving 100% polarization of cells, which is confirmed by our FACS results. This acknowledgment underscores the importance of exercising caution and precision in the interpretation of results. It highlights the necessity for a thorough understanding of the limitations inherent in the current methodology employed in this study and the advantage of using a machine learning model to accurately predict phenotypes for this study.

After confirming the degree of macrophage polarization, we next investigated the uptake of DNA-SWCNTs into the cells via confocal Raman microscopy. Macrophage cells that had been pre-polarized for 24 hours were treated with 1 mg-L^−1^ of GT_6_-SWCNTs for 30 minutes, followed by a thorough washing with 1x phosphate-buffered saline (PBS), and subsequent incubation in fresh media for an additional 30 minutes. After this incubation period, the cells were fixed and immersed in PBS for confocal Raman microscopy. Transmitted light and confocal Raman images were captured from individual cells at a magnification of 100x, aiming to analyze characteristic SWCNT Raman features such as the G-band and the D-band (figure 2a). The G-band, measured at 1585 cm^−1^,^48,18^ exhibits a linear correlation with SWCNT concentration (figure S5), while the D-band (1350 cm^−1^)^49^ to G-band ratio is indicative of the number of defects present on the SWCNT structure.^50^ Figure 2b represents the average integrated intensity of the G-band per region of interest (ROI). Here, an ROI defines a particular cell containing SWCNTs. The average integrated G-band intensity per ROI was found to be highest for M1 (42 counts), followed by M2 (23 counts), and then NM (13 counts). Figure S6, shows the full graphical representation of these cells.

**Figure 2.**
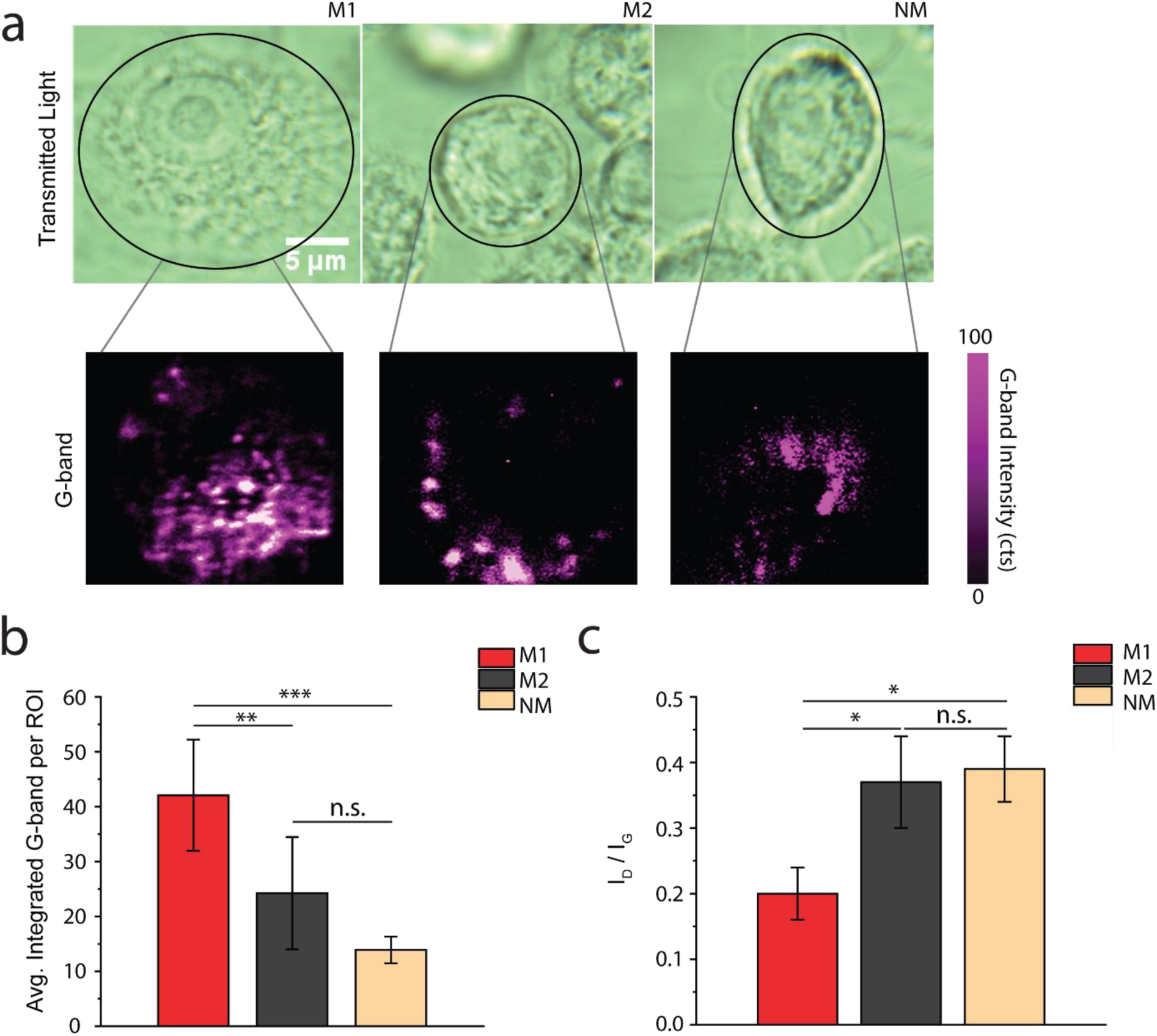
Raman microscopy characterization of cells containing DNA-SWCNTs. (a) Transmitted light images and magnified Raman G-band Images separated by cell phenotype. The scale bar on images is 5 µm. (b) Average integrated G-band intensity per ROI (c) D-band to G-band ratio showing clear differences in all cell phenotypes. Bars represent the average, and whiskers represent mean ± s.d. for each condition (*n* ≥ 4 cells per condition). Two-Sample t-test hypothesis testing analysis was performed between different samples and between different time points. (\*\*\**p* < 0.001, *\*\*p < 0.01* and \**p* < 0.05).

A statistical representation of the defect ratio (I_D_/I_G_) for DNA-SWCNTs within all macrophage phenotypes is presented (figure 2c). The data reveal that the defect ratio in M1 macrophages is significantly lower compared to the M2 and NM cells, and that there is no statistically significant difference between M2 and NM. A closer look at these observations suggests some selectivity in defective SWCNT uptake in M2 and NM cells. Our previous studies have shown that macrophages can induce defects and selectively internalize defective SWCNTs. This supports our hypothesis that M2 and NM cells selectively internalize SWCNTs with more defects, thus having a higher defect ratio.^51^ These results underline the intricate relationship between nanotube interaction and the internal cellular make-up, providing valuable insights into the nanomaterial cell dynamics.

Near-infrared hyperspectral fluorescence microscopy^15^ was performed on macrophage cells with internalized DNA-SWCNTs. All data was acquired as a function of DNA length. Three oligonucleotide sequences (GT_6_, GT_15_ and GT_30_) were employed in this study. DNA sequences composed of guanine-thymine (GT) repeat units were chosen for their widespread and comprehensive study in literature. Detailed Analysis of GT_15_ and GT_30_-SWCNTs can be found in figure S7-15.

Each cell type (M1, M2, or NM) was incubated with 1 mg-L^−1^ of GT_6-_SWCNTs for 30 minutes, followed by a wash with PBS, and subsequent incubation in fresh media for 30 minutes, 6 hours, or 24 hours before hyperspectral imaging. Throughout this period, changes in peak intensity and center wavelength were observed for four distinct emission bands in the NIR spectrum for each macrophage phenotype. From shortest to longest wavelength, these four emission bands are dominated by the (10,2), (9,4), (8,6), and (8,7)-SWCNT chiralities.^46^ Figure 3a shows the transmitted light and NIR images for all three macrophage phenotypes, M1, M2 and NM 6 hours post DNA-SWCNT dose. Visual differences in intensity can easily be observed with M1 being brighter than M2 and NM. Figure 3b shows the average broadband intensity for each cell type normalized by their respective areas. In principle, the NIR broadband intensity of M1 macrophages should ideally be significantly higher than that of M2 and naïve macrophages due to higher uptake in M1 as indicated by a higher G-band. However, it is important to note that the average cell size of M1 macrophages, as discussed earlier in figure 1b, is significantly larger in comparison to M2 and NM cells. When divided by the projected cell area in two-dimensions, this normalized broadband intensity turns out to be the smallest. The average area-normalized broadband intensity for M1 cells was observed to be a striking 72% less than M2. Furthermore, the average intensity for NM cells was observed to be approximately 45% less than that of M2 type macrophages. Nevertheless, it is crucial to emphasize that, whether normalized by cell area or not, the broadband intensity differs significantly across all cell phenotypes. This distinction suggests that the broadband intensity can serve as an identifiable feature for distinguishing cell phenotypes. Figure 3c shows averaged spectral data at the 6-hour time point, which reveals distinct differences and variations in normalized intensities and peak center wavelengths among M1, M2, and NM cells. Similar apparent trends and modulation in center wavelengths and peak intensities can be observed between these cell phenotypes throughout all time points (figure S16-18). To study identifiable spectral differences between M1, M2 and NM cells, we probed various NIR fluorescence parameters in more detail. Figure 3d is a statistical comparison of the peak intensity ratio for band 1 and 2. Statistically significant differences between all cell phenotypes is observed.

**Figure 3.**
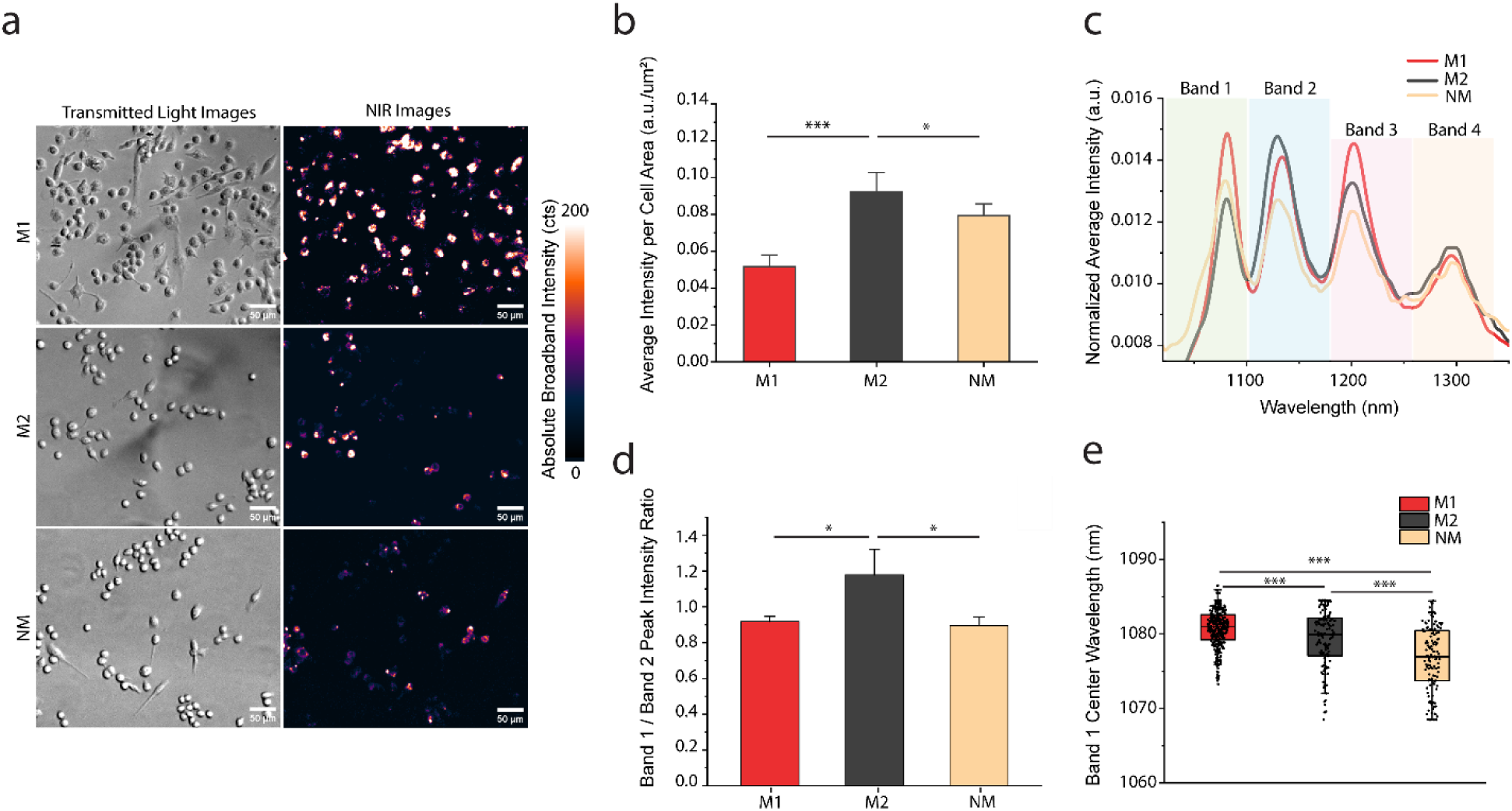
NIR fluorescence hyperspectral microscopy and identification of spectral features for GT_6_-SWCNTs in various macrophage phenotypes 6 hours after internalization. (a) Transmitted light and NIR fluorescence images (900-1600 nm) for all cell types M1, M2 and NM. (b) Bar graph showing average broadband intensity per cell area for all cell phenotypes. Bars represent the average, and whiskers represent mean ± s.d. for each condition (*n* ≥ 300 cells per condition). Two-Sample t-test hypothesis testing analysis was performed between different samples and between different time points. (\*\*\**p* < 0.001, and \**p* < 0.05). (c) Average NIR spectrum for GT_6_-SWCNTs within each cell phenotype. (d) Bar graph showing comparison of band peak intensity ratios for all phenotypes. Bars represent the average, and whiskers represent mean ± s.d. for each condition (*n* ≥ 300 cells per condition). Two-Sample t-test hypothesis testing analysis was performed between different samples and between different time points. (\**p* < 0.05). (e) Box and whisker diagram for band 1 center wavelength. Minimum of *n* ≥ 300 cells per condition were used. Boxes represent 25–75% of the data, horizontal lines represent medians, and whiskers represent mean ± s.d. Two-Sample t-test hypothesis testing analysis was performed (\*\*\**p* < 0.001).

We next fitted the average NIR spectra to Gaussian curves to extract peak intensity and center wavelength information for each of the identified SWCNT band (figure S19). The center-wavelength information is shown for the 6-hour timepoint in figures 3e, where it is apparent that both M1 and M2 type macrophages exhibit center wavelength modulations when compared to NM. M1 macrophages exhibited a 4 nm red-shift in band 1 center wavelength when compared to NM cells, whereas M2 macrophages exhibited a 3 nm red-shift. It is important to note that although the wavelength difference between M1 and M2 cells is just 1 nm, it is still statistically significant and thus can be used as one of the identifiable features in NIR macrophage phenotyping.

We speculate that the observed changes in intensity and center wavelength over 24 hours, as well as the differences in peak intensities and center wavelengths of different chiral species are likely attributed to variations in the lysosomal environment of these three distinct cell phenotypes.

Our previous work has demonstrated that DNA-SWCNTs undergo internalization into cells through energy dependent endosomal uptake and endo-lysosomal processing.^21,52^ It is also a known fact that the endosomal environment for different cell phenotypes, such as M1 and M2 macrophages, can significantly differ.^53^ For instance, M1 macrophages exhibit upregulation of NOX2, iNOS, SYNCRIP, TRAF6, AP-1 and certain Cathepsin species, while M2 macrophages show downregulation of NOX2, iNOS, SYNCRIP, and upregulation of Arginase, EGR2, SOD 1, and superoxide dismutase.^54,55,56^ The significant differences in the intra-endosomal environment of these cells contribute to a complicated interplay between SWCNTs and cellular biomolecules. This interplay leads to preferential binding of specific analytes to different SWCNT chiral species, thus influencing the SWCNT spectrum.

To confirm this hypothesis, we performed solution phase experiments to recapitulate the intracellular environments. Figure 4a shows the response of GT_6_-SWCNTs to physiologically relevant concentrations of Deoxyribonuclease II (DNase II), Activating Protein-1 (AP-1), and a mixture of both. DNase-II and AP-1 were selected to model the lysosomal environment of various macrophages due to phenotype-specific up or down regulation, as described previously. Similar to the differential phenotypic response as shown in figure 3, all the biomolecules exhibit slight changes in peak intensity and band specific center wavelength, which significantly differ from the GT_6_-SWCNT control sample. Notably, the individual biomolecule responses also significantly differ from the combined biomolecule response when carefully examining all NIR spectra. It is also important to note that like intracellular NIR features in figure 3 similar observations in center wavelength red-shifting and intensity changes are observed for the SWCNT bands between control, DNase-II and AP-1 samples. Interestingly, the protein-induced spectral modulations are time-dependent (figure 4c). Major shifts in SWCNT center wavelengths and peak intensities were observed for AP-1 solutions at timepoints of 1 hour or 6 hours after mixing. Similar time-dependent spectral modulations were observed for DNase-II and the biomolecule mixture as shown in figure S20. These differences in SWCNT sensor response over time, specifically peak shifting and broadening can also be attributed to protein-induced aggregation within the samples^57^. The differential response of the DNA-SWNCTs under varying pH conditions is shown in figure 4b. Finally, figure 4c illustrates variations in the peak intensity ratio between band 3 and 4 across all model biomolecules, as depicted by the bar graph.

**Figure 4.**
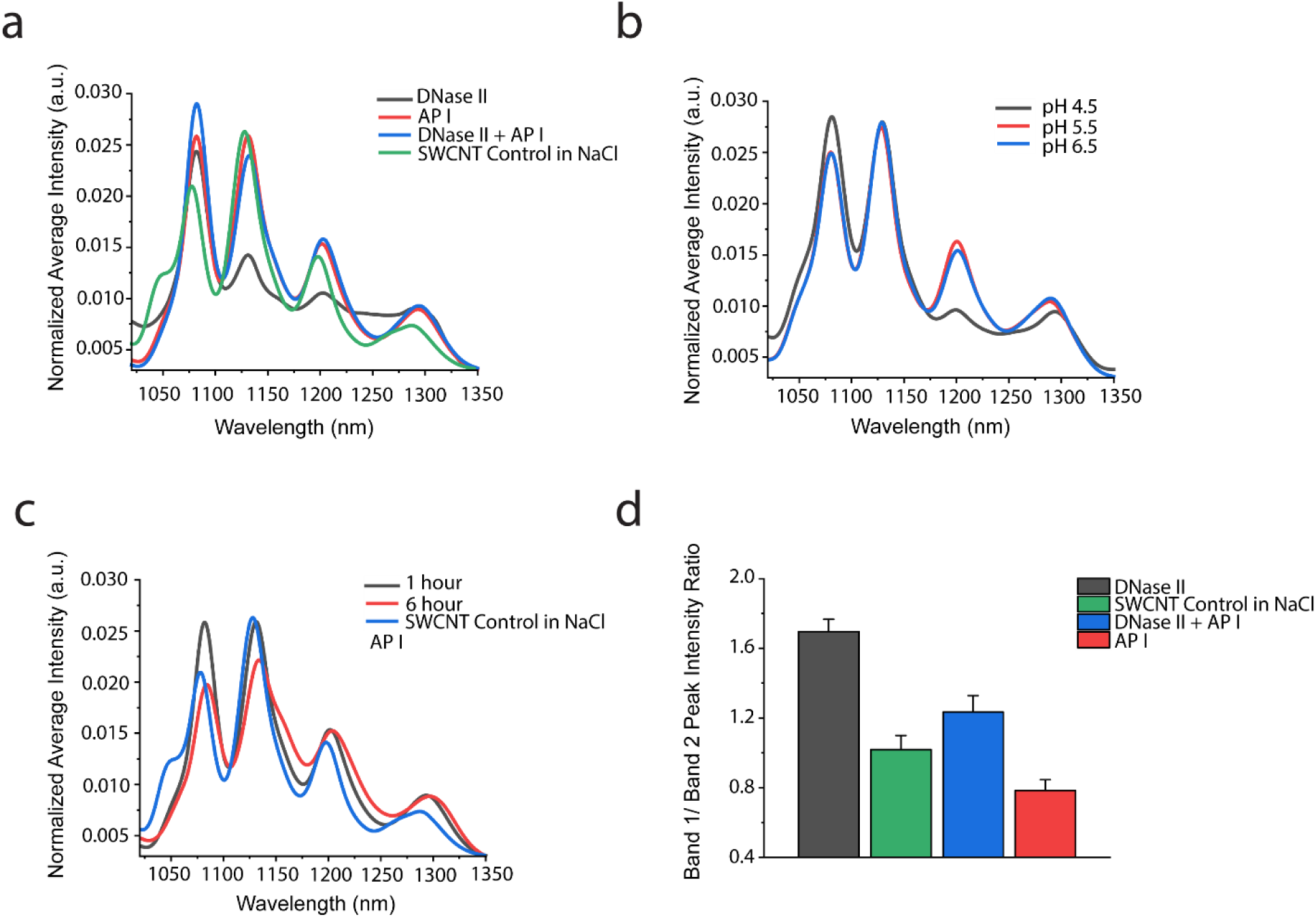
Solution phase experiments recapitulating the NIR fluorescence response of GT_6_-SWCNTs to the endo-lysosomal environment. (a) Spectral response to DNase-II and AP-1. (b) Spectral response to varying pH environments. (c) Time-dependent response to both DNase-II and AP-1. (d) Bar graph showing comparison of band peak intensity ratios for all phenotypes.

These solution phase experiments provide supporting evidence for the hypothesis that differences in the NIR spectrum of DNA-SWCNTs can indeed be caused by changes in the varying endo-lysosomal environment within distinct macrophage phenotypes.

**Figure 5.**
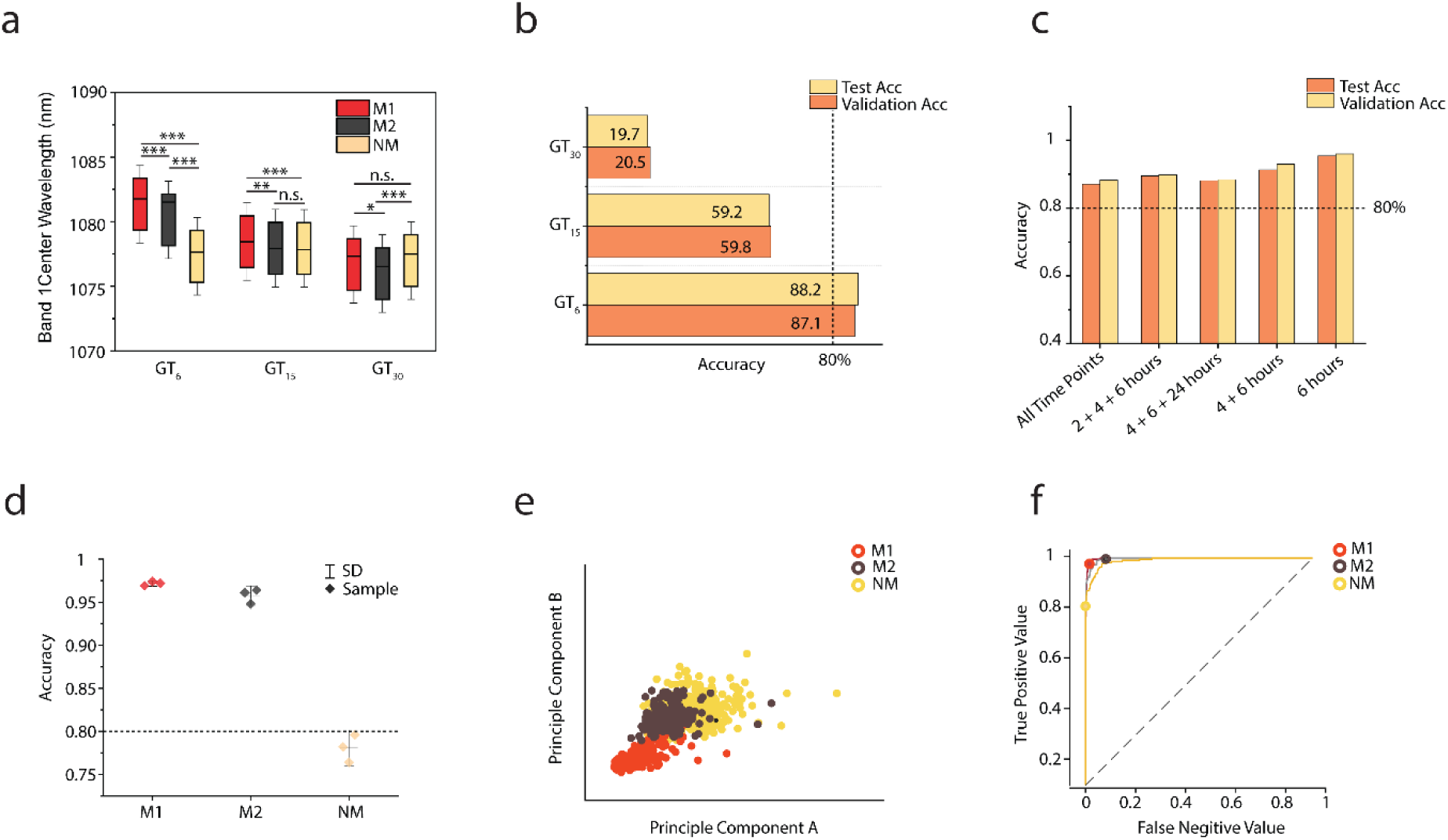
Machine Learning for Phenotype Identification. (a) Band 1 center wavelength feature comparison between three different DNA-SWCNTs, i.e., GT_6_, GT_15_ and GT_30_. (b) Bar graph indicating accuracy of test and validation data for all tested DNA-SWCNTs. (c) Bar graph indicating accuracy of test and trained data for GT_6_-SWCNT evaluated for different time point data sets. (d) Interval plot indicating accuracy of predicting each macrophage phenotype. (e) Principal component A and B cluster plot showing each cell type cluster. (f) ROC curve for each macrophage phenotype indicating true positive and false negative values.

The array of changes observed in the DNA-SWCNT spectra across different cell phenotypes and over time reflects the complex and dynamic nature of nanomaterial-cell interactions within distinct endo-lysosomal environments. This insight into the intricate molecular interactions can be used as an effective tool to differentiate between cell phenotypes. Utilizing all the NIR fluorescence features that exhibited significant differences between M1, M2, and NM cells, a machine learning (ML) model was developed for the identification and quantification of unknown macrophage samples into either the M1, M2 or NM cell category. We selected Support Vector Machine (SVM) as the ML model for this purpose. SVM is a supervised learning algorithm designed for solving complex classification, regression, and outlier detection problems.^58^ It achieves this by performing optimal data transformations that delineate boundaries between data points based on predefined class labels or outputs. SVM modeling has proven effective in various applications and has recently gained momentum within healthcare and medicine.^59,60,61^ This method has previously been employed for the detection of various diseases such as Diabetes,^62^ Alzheimer’s,^63,64^ Psoriasis,^65^ Hepatitis,^66^ and more.^67,68^

Utilizing this SVM ML model, we examined the ability of DNA-SWCNTs to identify differing cell phenotypes as a function of DNA length. In figure 4a, one identifiable feature, i.e. band 1 center wavelength, is shown as a function of DNA length and macrophage phenotype. Upon statistical analysis, DNA-SWCNTs composed of the shortest DNA sequence, i.e. the GT_6_, exhibited the largest statistically significant modulations. Additional comparisons are found in figure S21 and S22. With this DNA length-dependence in mind, we trained individual SVM models based on NIR fluorescence spectra of DNA-SWCNTs within macrophages of known phenotype. For GT_6_-SWCNTs, although 101 NIR features were input into the model, a principal component analysis and feature importance score sorting using an ANOVA analysis were performed (figure S23) and only 42 significantly identifiable features explaining at least a 95% variance were selected. Figure 4b shows a bar graph demonstrating that for both validation and test data, GT_6_-SWCNTs achieved the highest accuracy at about 87%. GT_15_ exhibited moderate accuracy, reaching approximately 59%, while GT_30_ significantly underperformed with an accuracy of only 19%. The SVM ML accuracies were determined as a function of timepoint (figure 4c). All models trained with GT_6_ data consistently achieved accuracies greater than 87%. Notably, three of the GT_6_ models surpassed 90% accuracy, and the model trained with data from the 6-hour time point exhibited the highest accuracy at approximately 95%. When delineated by cell phenotype, the model demonstrates the highest accuracy for predicting M1 and M2 macrophages, with 98% and 96% accuracies, respectively. The accuracy for NM cells was comparatively lower at roughly 78%. The lower accuracy of NM cells is attributed to a significant overlap of principal component analysis features between M2 and naïve macrophages, as depicted in figure 4e. Figure 4f shows a receiver operating curve (ROC), further affirming the high predictive power of the model. The ROC curve is a valuable tool for assessing the tradeoff between sensitivity and specificity in classification models.

The higher predictive accuracy, especially for M1 and M2 macrophages, and the corresponding ROC curve, contribute to the overall confidence in the robustness and reliability of the GT_6_ SVM model for identifying and distinguishing between different macrophage phenotypes. Furthermore, these results also underline the consistency of the GT_6_-SWCNT model across various time points, with particularly high accuracy rates observed. The superior performance of the model trained with data from the 6-hour time point suggests that this specific time point provides highly discriminatory features for accurately identifying cell phenotypes. It is imperative to note that number of data points to train data were kept constant so avoid any issues arising from over-fitting or under-fitting of data. It is also important to note that previous studies have shown this 6-hour time scale is very significant for intracellular studies. Machine learning data for other timepoints can be found in supplemental information, figure S24 and S25. Overall, the findings emphasize the potential of GT_6_-SWCNT data in developing accurate and reliable machine learning models for cell phenotype identification.

For future *in vivo* applications of the spectral fingerprinting approach, we demonstrate feasibility with the use of a custom-built NIR probe spectrophotometer.^69^ A schematic of the probe spectrophotometer is shown in figure 6a. The setup incorporates a 730 nm laser passing through a two-way fiber optic cable and illuminating a petri dish of M1, M2, or NM cells containing DNA-SWCNTs. The fluorescence emission is directed back into the fiber optic probe, which routes it into a spectrometer connected to a 1D InGaAs detector, providing an output in the form of a NIR spectrum. Figures 6b and 6c represent the normalized output from the probe spectrophotometer at 0.5 hours and 2 hours after a 30-minute incubation with 5 mg-L^−1^ GT_6_-SWCNTs. Evident intensity changes between the two band species can be observed. Figure 6d shows the differential peak intensity ratio comparisons for all phenotypes at 0.5 hours post SWCNT dose. Significant differences in intensity ratios observed are features that can easily enable the identification of macrophage phenotypes with considerable accuracy.

**Figure 6.**
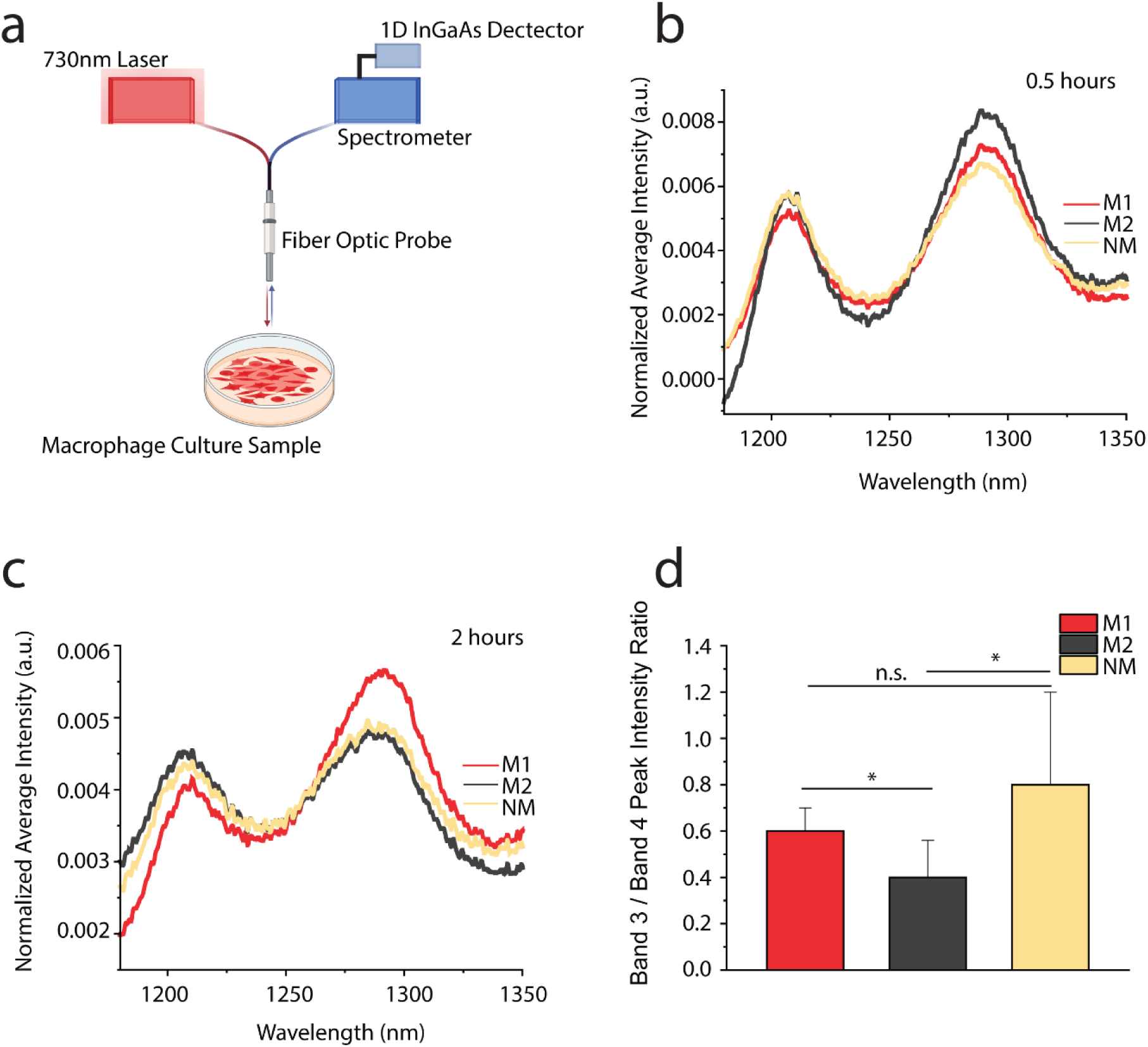
Simulating *in vivo* applicability of the spectral fingerprinting approach. (a) Schematic diagram showing fiber optic NIR probe spectrometer setup for sample data collection. (b) GT_6_-SWCNT sensor probe response at 0.5 hours. (c) GT_6_-SWCNT sensor probe response at 2 hours. (d) Bar graph showing comparison of band peak intensity ratios for all phenotypes. Bars represent the average, and whiskers represent mean ± s.d. for each condition (*n* ≥ 5×10^5^ cell/ml per petri dish). Two-Sample t-test hypothesis testing analysis was performed between different samples and between different time points. (\**p* < 0.05).

To evaluate the use of DNA-SWCNT-based cell phenotyping in a more clinically relevant application, i.e. in primary cells, the same type of NIR spectral analysis was performed with bone marrow-derived macrophages (BMDMs) isolated using bone marrow from femoral and tibial bones of eight- to ten-week-old mice. Figure 7a and figure S26 is a visual representation of all the cell phenotypes and clearly shows the apparent differences in NIR brightness among all three cell types. Interestingly, like previous figures, significant band center wavelength and intensity modulations are observed among M1, M2 and naïve BMDM samples through time as shown by figure 7b and 7c. Similarly, a closer look at these features indicates statistically significant differences in the band peak intensity ratios when comparing all three cell phenotypes as shown in figure 7d. More distinguishable NIR features can be found in the supplemental information, figure S27 and S28. Together, these distinct differences demonstrate the applicability of our sensor platform towards *in vivo* applications, offering the potential for real-time monitoring and analysis of cellular responses and comparison of different cell phenotypes present within living organisms.

**Figure 7.**
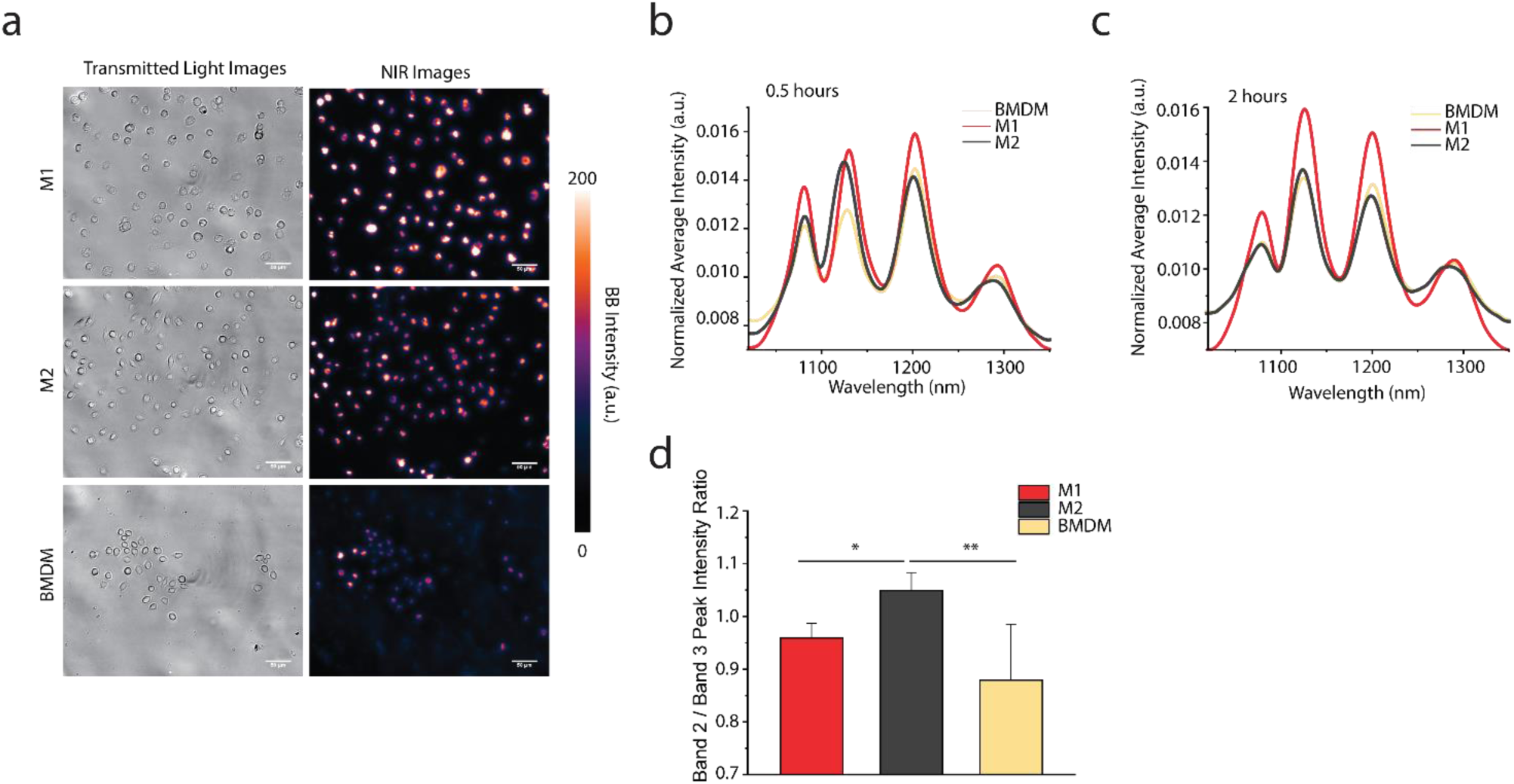
Simulating applicability of the spectral fingerprinting approach in primary BMDM cells. (a) Transmitted light and NIR fluorescence images for all cell types M1, M2 and naïve BMDM at 0.5 hours. (b) Normalized average intensity for all cell phenotypes at 0.5 hours. (c) Normalized average intensity for all cell phenotypes at 2 hours. (d) Bar graph showing comparison of band peak intensity ratios for all phenotypes. Bars represent the average, and whiskers represent mean ± s.d. for each condition (*n* ≥ 300). Two-Sample t-test hypothesis testing analysis was performed between different samples and between different time points. (\*\**p* < 0.01 and \**p* < 0.05).

To quantify potential deleterious effects on cell health as a function of polarization state and DNA-SWCNT concentration, two distinct methods, xCELLigence and Annexin V/PI assay, were employed to assess the response of each cell phenotype. The first method involved real time monitoring of cell proliferation using an xCELLigence Real-Time Cell Analysis system. This system measures electrical impedance across integrated micro-electrodes embedded in the bottom of specialized 16-well tissue culture E-plates. The impedance measurement is represented as a cell index value, which can be directly correlated to various cellular characteristics, including cell growth, viability, adhesion, activity, and morphology.^70^ Figure 8a illustrates the normalized cell index of RAW 264.7 NM cells dosed with 0.1, 1, or 10 mg-L^−1^ GT_6_-SWCNTs. The data reveal that the highest cell proliferation occurred at a DNA-SWCNT dose of 1 mg-L^−1^, while a dose of 10 mg-L^−1^ considerably decreased cell viability. Interestingly, higher DNA-SWCNT concentrations resulted in greater rates of cell proliferation, activity, and viability of M1 macrophages, with 10 mgL^−1^ showing the highest response (figure 8b). The response of M2 macrophages mirrors that of NM cells, with maximum growth and proliferation observed at a dose of 1 mg-L^−1^ and a minimum growth rate observed at 10 mg-L^−1^ (figure 8c).

**Figure 8.**
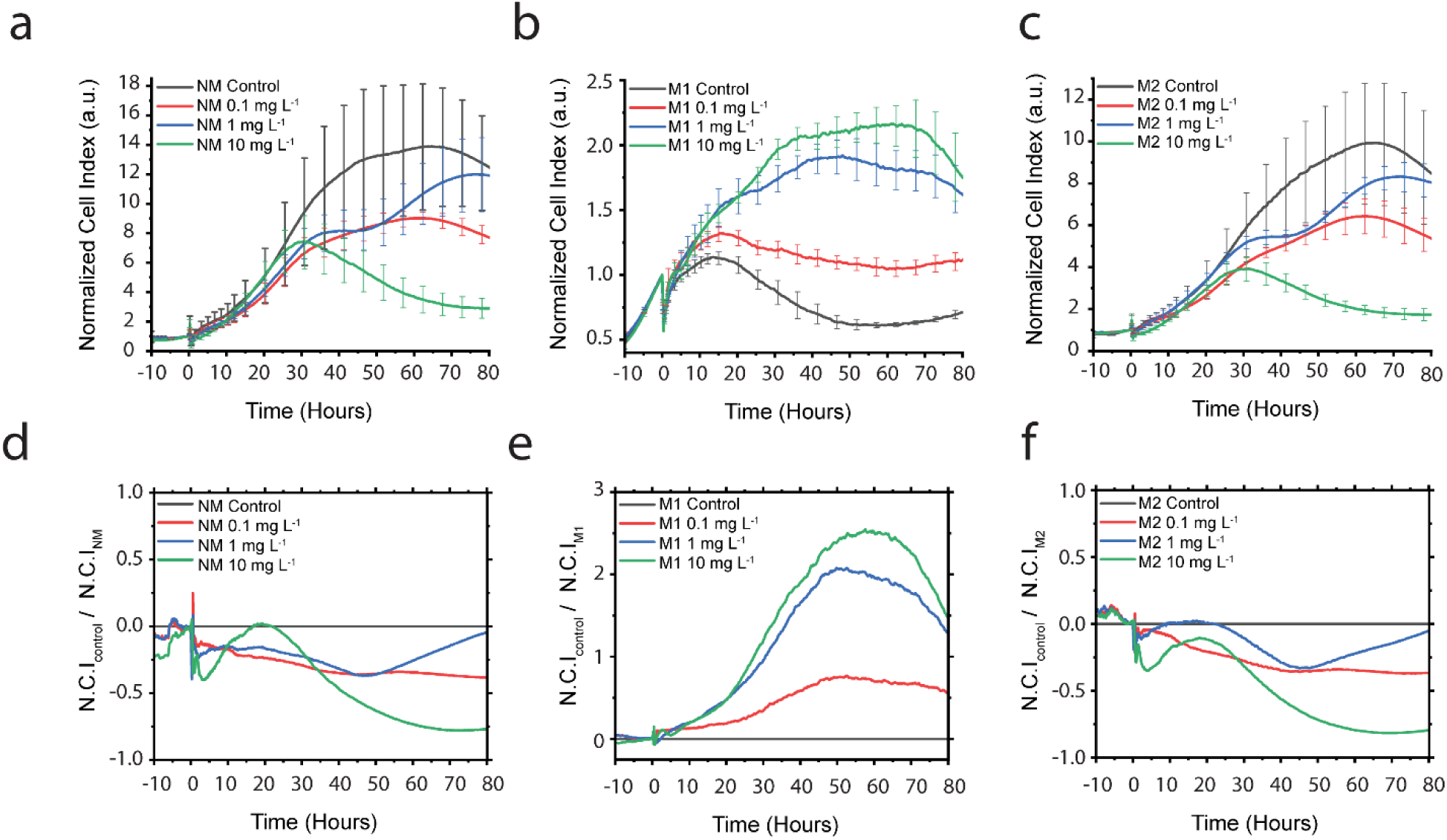
Real-time cell proliferation and cytotoxicity of RAW 264.7 macrophages exposed to DNA-SWCNTs. Cell proliferation curves for (a) naïve macrophages, (b) M1 macrophages, and (c) M2 macrophages over a continuous period of 3 days. Cell index normalized to the no-SWCNT control cell index for (d) naïve macrophages, (e) M1 macrophages, (f) M2 macrophages, as a relative measure of cell activity.

By normalizing the cell index of each macrophage phenotype to the cell index of its respective control, a measure for relative cellular activity over time was determined (figures 8d-f). Both M2 and NM cells exhibit an identical response, with their activity lower than that of their respective controls. Upon closer examination of M1 macrophages, it is observed that all M1 macrophages dosed with DNA-SWCNT outperform the control in terms of cellular activity, with the highest nanotube dose demonstrating the highest activity. The enhanced activity of M1 macrophages is attributed to their higher uptake of DNA-SWCNTs. We speculate, due to their pro-inflammatory nature, M1 macrophages tend to proliferate more when internalizing more DNA-SWCNTs. This observation suggests that the presence of higher concentration of SWCNTs may stimulate the activity and growth of M1 macrophages, providing valuable insights into the complex interplay between nanomaterials and immune cell responses.

A second method for monitoring cell health was an apoptosis-necrosis assay, performed to investigate the effects of DNA-SWCNT concentration on different macrophage phenotypes as shown in figure S4. Results did not show any major cell necrosis in any samples except M1 cells. This was due to scrapping the M1 cells from the petri dish surface, which has been discussed previously.

**Figure 9.**
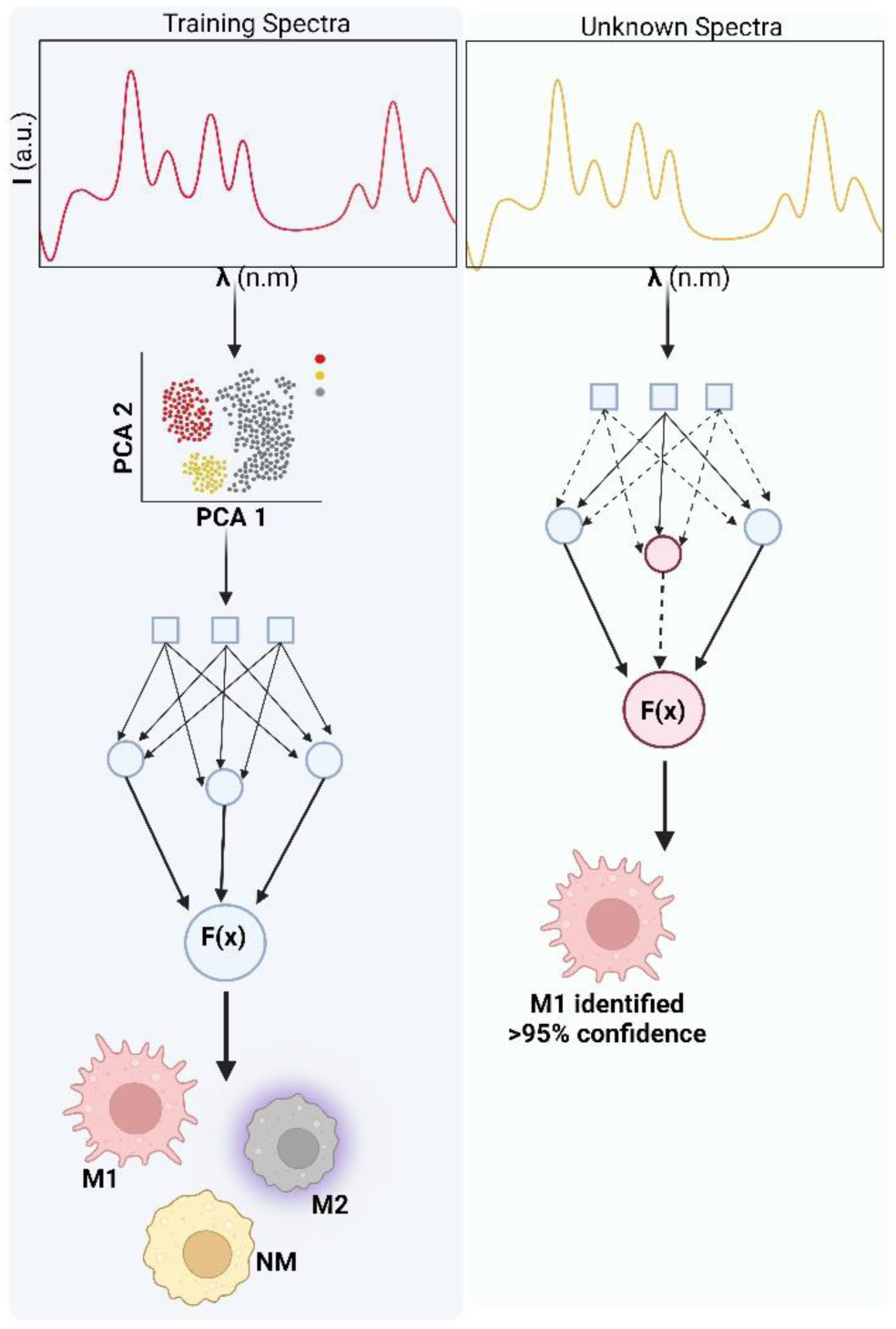
Concluding schematic. A simplified schematics for the accurate identification of immune cell phenotypes using a support vector machine learning model.

## Conclusion

In conclusion, this comprehensive study employs a multifaceted approach to unravel the intricate interactions between distinct macrophage phenotypes and DNA-SWCNTs. A crucial finding highlights the paramount importance of selecting an appropriate DNA sequence length for accurate cell phenotype identification based on NIR fluorescence data, with an emphasis on efficacy of shorter sequences. Moreover, the study affirms the potency of SWCNT-based spectral fingerprinting, particularly when coupled with machine learning, as an invaluable tool for precisely categorizing cellular states based on complex spectral data. This innovative approach holds great promise in advancing our understanding of dynamic cellular processes, disease states, disease progression, and response to stimuli *in vivo*. The platform’s sensitivity to minute changes in the NIR spectrum positions it as a valuable tool for researchers and clinicians. It facilitates the real-time monitoring of cellular behavior within living systems, offering insights that extend to both fundamental research and clinical applications. The implications of this research pave the way for future developments in nanotechnology and biomedicine, bridging the gap between spectral analysis and cellular dynamics for enhanced diagnostic and therapeutic interventions.

## Methods and Materials

### DNA-SWCNT Sample Preparation

To create monodispersed ssDNA wrapped SWCNTs (ssDNA–SWCNTs), 1 mg of as-synthesized SWCNT powder was added to 2 mg of (GT)_xx_ oligonucleotide (Integrated DNA Technologies) 1 mL of 0.1 M NaCl (Sigma-Aldrich). Each sample was ultrasonicated using a 1/8′′ tapered microtip for 30 min at 40% amplitude in an ice bath (Sonics Vibracell VCX-130; Sonics and Materials). The resulting suspensions were ultra-centrifuged (Beckman Optima MAX-XP) for 30 min at 250,000 *g* and 4°C, and the top ~80% of the supernatant was collected.

### Near-Infrared Fluorescence Microscopy

A hyperspectral NIR fluorescence microscope, similar to a previously detailed system, was used to obtain all hyperspectral fluorescence data. A 660 nm excitation laser source was reflected onto the sample stage of an Olympus IX-73 inverted microscope equipped with a LCPlan N, 20x/0.45 IR objective by Olympus, U.S.A. The resulting fluorescence emission was passed through a volume Bragg grating and collecting with a 2D InGaAs array detector by Photon Etc., (Montreal, Canada) to generate spectral image stacks. Live cell samples were mounted on a stage top incubator by Okolab, to maintain 37°C and 5% CO_2_ cell culture conditions throughout the imaging procedure. All hyperspectral cubes, Fluorescence images and transmitted light images were corrected and processed in MATLAB.

### Confocal Raman Microscopy

All Raman data was acquired using an inverted WiTec Alpha300R confocal Raman microscope (WiTec, Germany) equipped with a ZEISS Epiplan-NEOFLUAR 100x/1.3 Oil Pol, Oil immersion, objective, a 785nm laser (20 mW output measured at the sample), and a UHTS 300 spectrograph (300 lines/mm grating) coupled with an Andor DR32400 CCD detector (−61C, 1650 × 200 pixels). Singular cell areas were scanned, and spectra were obtained in 1 × 1 um intervals using a 0.2s integration time per spectrum to construct hyperspectral images of individual cells. Background subtraction and cosmic ray removal were performed using a polynomial function in WiTec Project 5.2 software. Hyperspectral data was extracted and processed using custom codes written with MATLAB.

### Cell Culture

RAW 264.7 TIB-71 cell line from ATCC (Manassas, VA, USA) was cultured under standard incubation conditions at 37C and 5% CO2. D-10 cell culture media containing sterile filtered high-glucose DMEM with 10% heat inactivated FBS, 2.5% HEPES, 1% L-glutamine, 1% penicillin/streptomycin, and 0.2% amphotericin B (all by Gibco) was used for cell culture. 0.5ng/ml of cytokines and signaling molecules were added to media during each experiment to avoid cells losing their polarization states.

### Sample Preparation for Optical Microscopy

For all 20x *in vitro* NIR fluorescence imaging experiments the cells were plated, in triplicate, at an initial concentration 5.26 × 10^4^ cells/cm^2^ on 35 mm glass-bottom microwell dishes (MatTek) and allowed to culture overnight. To dose the cells with nanotube samples, the culture media was removed and replaced with 1 mg-L^−1^ of either purified-SWCNTs or unpurified-SWCNTs diluted in D10 cell culture media and incubated for 30 minutes to allow cell internalization. The SWCNT-containing media was then removed, the cells were washed twice with sterile PBS (Gibco) followed by the addition of fresh media. All time points were defined with respect to this step. For the 100x Confocal Raman microscopy experiments, the same cell plating and SWCNT dosing procedures were followed as highlighted previously; however, the cells were fixed with paraformaldehyde (Electron Microscopy Sciences). Cell fixation was performed with 4% PFA in PBS for 15 minutes, after which the cells were rinsed three times and covered with PBS to retain an aqueous environment during imaging.

### Cell Viability Assay

RAW 264.7 macrophage cells were plated on 35 mm glass-bottom microwell dishes (MatTek) and allowed to culture overnight at an initial seeding density of 5.26 × 10^4^ cells/cm^2^. The following day, the medium was replaced with 1 mg/L of either purified-SWCNTs or unpurified-SWCNTs diluted in media and incubated for an additional 24 h. After 24 h, the cells were collected from the dishes and stained with Annexin V and propidium iodide (Dead Cell Apoptosis Kit V13242, Invitrogen) following the manufacturer’s protocol. Fluorescence images of the stained cells were acquired by using a Cellometer Vision CBA image cytometer (Nexcelom Bioscience), and images were analyzed by using ImageJ and custom MATLAB codes. For each cell condition, a control dish was plated without SWCNT addition to create the gates on the Annexin V and propidium iodide axes of the histograms.

### Bone Marrow Cell Isolation

To generate BMDMs, the femoral and tibial bones of 8-10-week-old mice were flushed with RPMI media (Thermo Fisher) and the bone marrow suspension was passed through a 70 mm cell strainer. Bone marrow cells were cultured in T75 non-tissue-culture flasks with 10 mL of RPMI medium (Thermo Fisher) supplemented with 10% heat-inactivated fetal bovine serum, 2 mM L-glutamine, 25 mM HEPES, and 20% L929 conditioned medium and kept in a humidified incubator at 37C with 5% CO2. An additional 5mL of media was added to plates on day 3 of differentiation. After 7-10 days of differentiation, the loosely adherent cells were harvested by gentle washing with 2 mM EDTA (Thermo Fisher) in PBS (Thermo Fisher), cells were seeded in 12-well plates at a density of 3 × 10^5^per mL, were pooled and used as the starting source of cells for most experiments. BMDMs were seeded overnight in 12-well plates (2.5×106/well; 1 ml culture medium before infection).

### Macrophage Differentiation

To induce differentiation of macrophages into M1 and M2 phenotypes, a combination of signaling molecules were added. M1 were obtained by exposing the cells to a dose containing 0.5ng/ml of lipopolysaccharide (LPS) and interferon-gamma (IFN-ɣ). LPS, derived from the outer membrane of gram-negative bacteria, and IFN-ɣ, a pro-inflammatory cytokine, collectively drive cell polarization towards the M1 phenotype. Conversely, to obtain M2, a separate set of signaling molecules is used. A dose of 0.5 ng/ml of interleukin-4 (IL-4) and interleukin-10 (IL-10) is introduced simultaneously. IL-4 and IL-10 cytokines are both associated with anti-inflammatory responses, which contribute equally to the polarization of macrophages towards the M2 phenotype. Cells were incubated with the cytokine mixtures overnight, for 24-hours, to achieve maximum cell polarization and were later evaluated for characteristic differences among each cell phenotype, before proceeding with any experimentation.

### Real-Time Near-Infrared Fluorescence Spectroscopy for In-vivo simulation

Petri dishes were seeded with cells to achieve a density of 5 × 10^5^ cells. Consequently, cells were dosed with respective cytokines to polarize them into M1 and M2 macrophages. After 24 hours, 5mg L^−1^ of GT6-SWCNTs were added to cells, after the 30-minute incubation period, cells were washed with 1x PBS and replenished with fresh media. NIR fluorescence spectra were acquired from each sample at 0.5 and 2 hours. Individual NIR fluorescence spectra from cell samples were obtained using a custom-built preclinical fiber optics probe spectroscopy system described in previous studies. A custom MATLAB code was used to perform background subtraction and post analysis on acquired fluorescence data.

### Macrophage Identification

To discern M1 macrophage characteristics, PE and PerCP-conjugated anti-mouse 1-A/I-E antibodies (BioLegend) were utilized. These antibodies were specifically employed for the precise detection of M1 macrophage markers, including CD4 CD3/TR. These selected markers play a pivotal role in identifying distinctive characteristics associated with M1 macrophages using flow cytometry. Subsequently, M2 macrophages were also identified. In this regard, M2-specific markers, CD206 and CD163, were targeted for detection using PE-Dazzle 594 (BioLegend). An Anti-CD16/CD32 Fc receptor blocking antibody was to eliminate non-specific receptor binding. This meticulous approach allowed for the comprehensive identification of M1 and M2 macrophage phenotypes post cytokine addition.

### FACS Flow Cytometry

Cells were scrapped off from petri-dishes and resuspended in FACS running buffer (PBS + 0.5-1% BSA) at a density of 5 × 10^6^ cells/ml. Staining was done at 4°C. 100 µl of cells were added to centrifuge tubes followed by 100µl of Fc blocking antibody (1:50 ratio in FACS running buffer). The cells were incubated for 20 minutes followed by centrifugation at 1500 rpm for 5 minutes at 4°C. Supernatant was discarded and the cells were further incubated for 30 minutes with marker specific antibodies (0.1 µg/ml) in 100 µl of FACS buffer, in the dark. The cells were then washed by centrifuging at 1500 rpm for 5 minutes. Washing was repeated 3 times to remove any unbound markers. The cells were then resuspended in 200µl of FACS buffer for flow cytometry.

### Label-Free Cell Proliferation and Adherence Monitoring

Adherence and proliferation were measured with an xCELLigence real-time cell analysis instrument from Agilent. For baseline impedance measurements of the wells, 140µl of cell media was added to each of the 16 wells in the E-plate. 50µl of cells diluted in media was added to each well to reach a final concentration of 2 × 10^5^ cells/well. The cells were allowed to adhere to the plates for 30 minutes in a cell culture hood to allow for an evenly distributed initial seeding of cells over the electrodes. After 30 minutes, the E-plates were placed into the xCELLigence system and data acquisition occurred every 15 minutes. Plates were incubated for 24 hours. Subsequently, respective cytokines were added, and the cells were incubated for an additional 24 hours to polarize macrophages into M1 or M2 phenotypes. The cells were then dosed with three different concentrations of (GT)_6_-DNA-SWCNTs: 0.1, 1, and 10 mg/L, in separate wells.

### Machine Learning

A custom-made MATLAB graphical user interface was used to format data for machine learning analysis. MATLAB classification learner was used to train data with the support vector machine model on the formatted data.

### Statistical Analysis

OriginPro 2022b was used to perform all statistical analyses. All data either met assumptions of statistical tests performed (i.e., equal variances, normality, etc.) or were transformed to meet assumptions before any statistical analysis was caried out. Statistical significance was analyzed using Two-Sample t-test or one way ANOVA where appropriate. Testing of multiple hypotheses was accounted for by performing one-way ANOVA with Tukey’s posthoc test.

## Supporting Information

The Supporting Information is available free of charge at:

Experimental methods and materials, additional characterization spectra, images, and analysis.

## Acknowledgements

This work was supported by the National Science Foundation (CAREER Award #1844536 and #2231621) and the University of Rhode Island College of Engineering. The confocal Raman data were acquired at the RI Consortium for Nanoscience and Nanotechnology, a URI College of Engineering core facility partially funded by the National Science Foundation EPSCoR, Cooperative Agreement #OIA-1655221. Research was made possible by the use of equipment available through the Rhode Island Institutional Development Award (IDeA) Network of Biomedical Research Excellence from the National Institute of General Medical Sciences of the National Institutes of Health under grant #P20GM103430 through the Centralized Research Core facility. Schematics were created using BioRender.com software.

