## Supplemental Information for "Machine Learning Assisted Spectral Fingerprinting for Immune Cell Phenotyping"

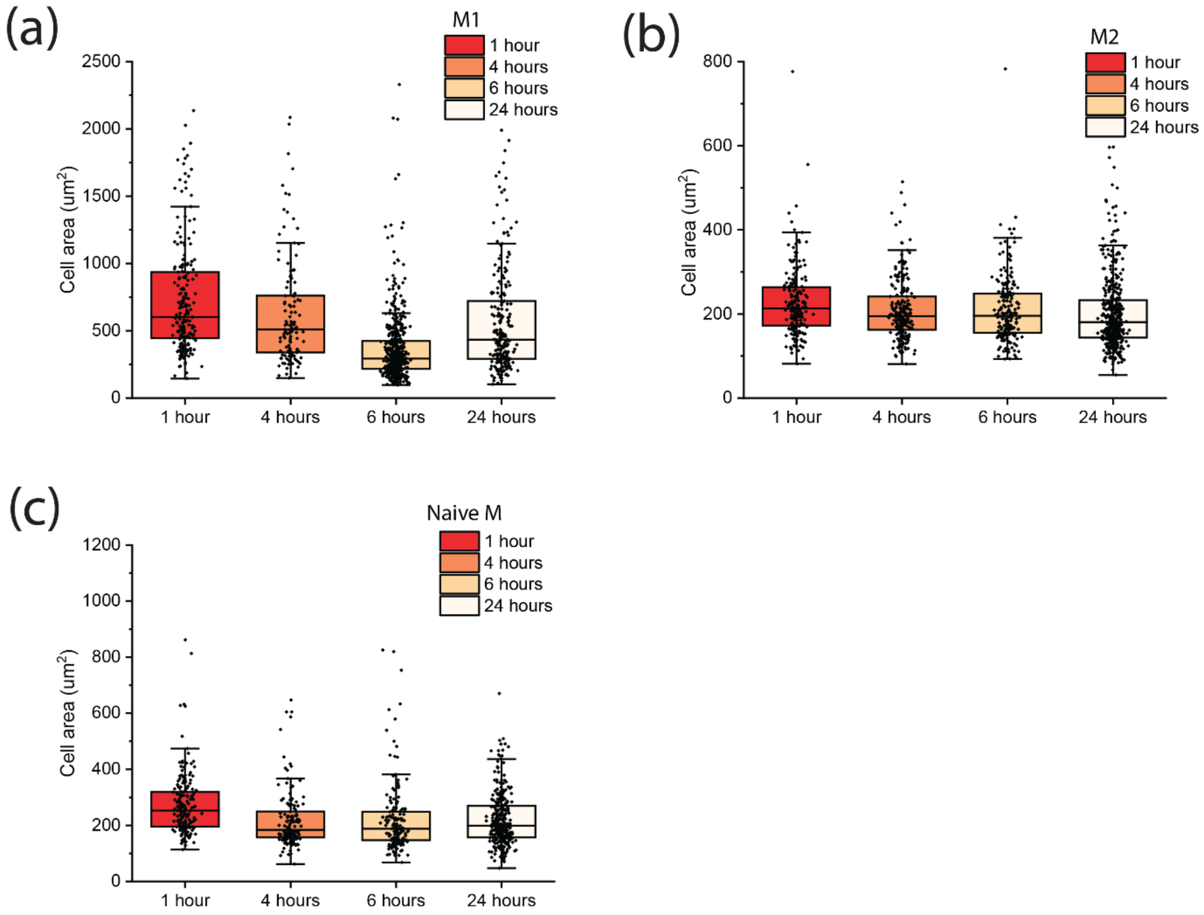


**Figure S1.** Size distribution analysis across macrophage phenotypes. Box and whisker plot showing size distribution of macrophage phenotypes (a) M1 (b) M2 (c) Naïve Macrophages taken at four different time points i.e. 1 hour, 4 hours, 6 hours, and 24 hours post polarization. Each ROI in the box and whisker plot is a single cell. Minimum of $n \geq 300$ cells per condition were used. Boxes represent 25–75% of the data, horizontal lines represent medians, and whiskers represent mean ± s.d.

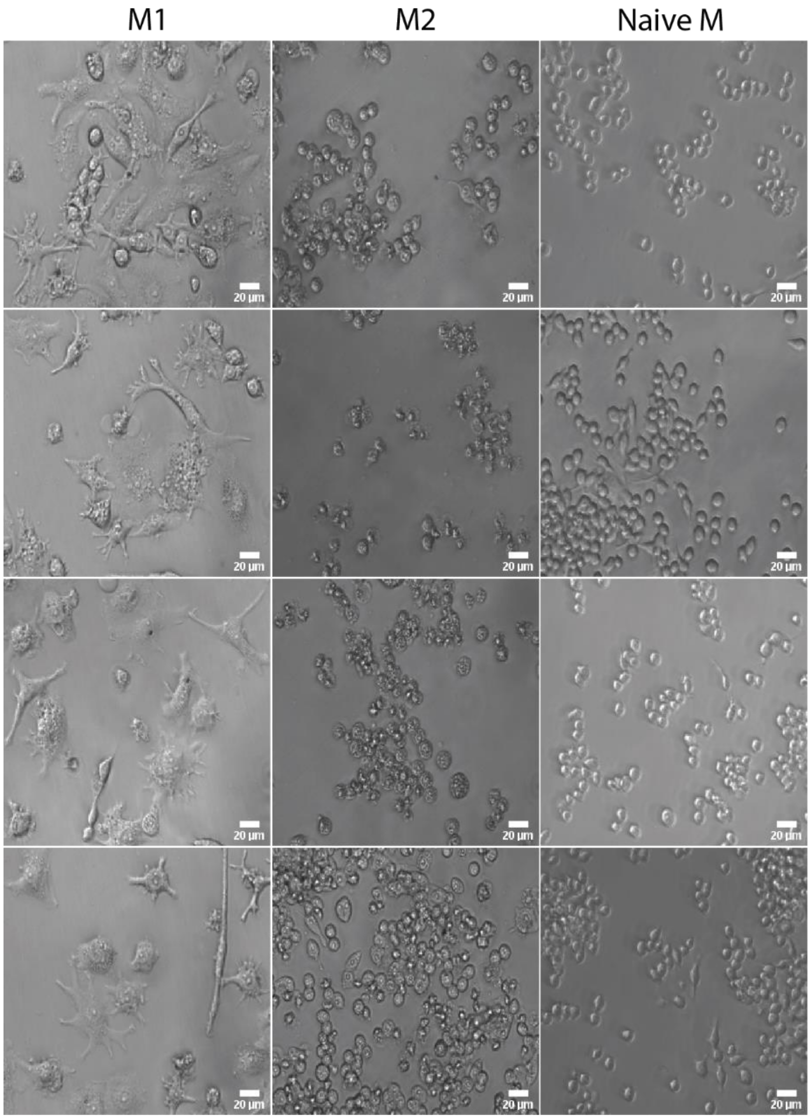


**Figure S2.** Transmitted Light Images of M1, M2 and Naïve Macrophages taken at 20x magnification. The scale bar for each image is 20μm.

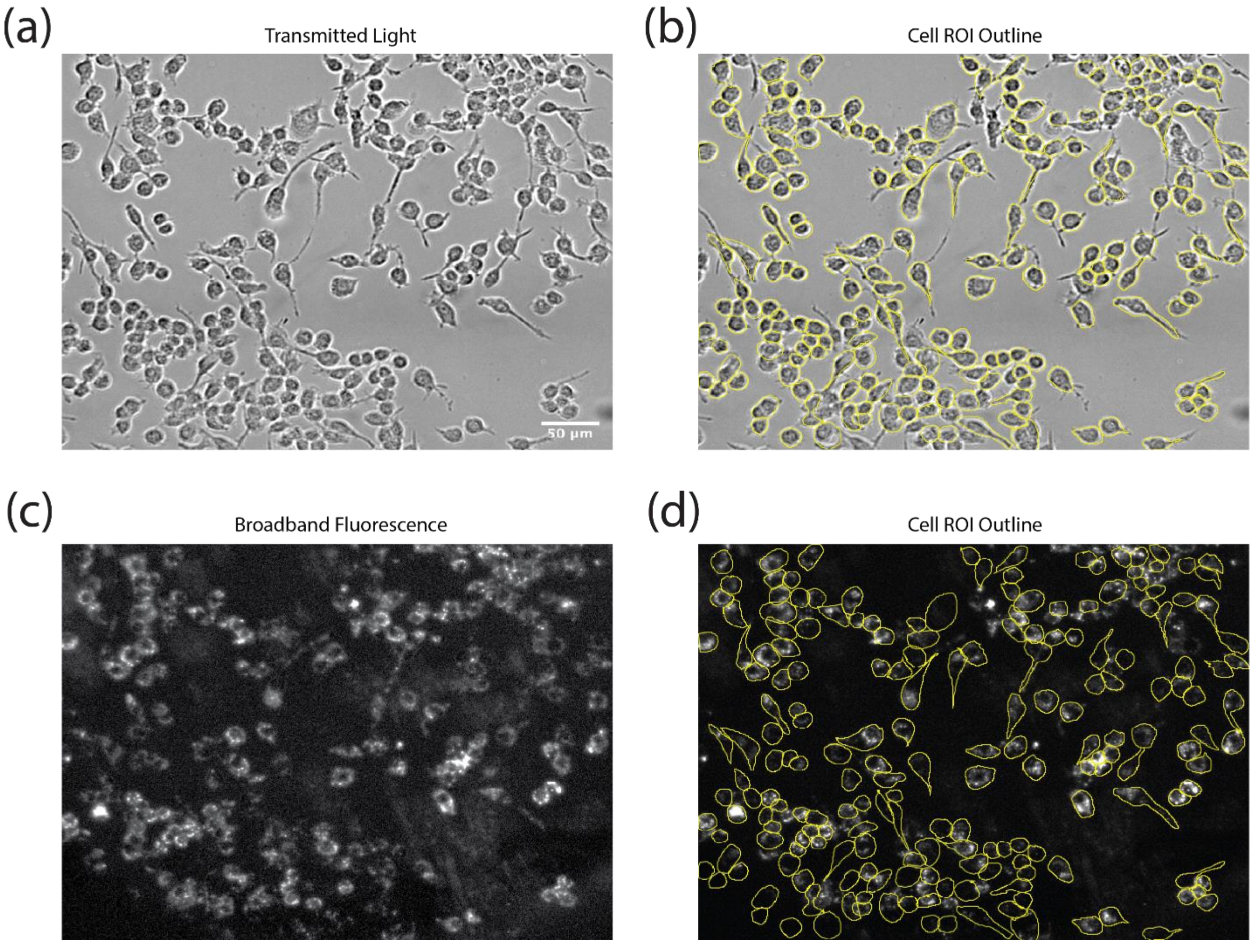


**Figure S3.** Sample images for single cell region of interest (ROI) analysis. (a) Transmitted light image and (b) broadband NIR fluorescence image of RAW 264.7 cells dosed with 1 mg $L^{-1}$ $GT_6$-SWCNTs. Right panels overlay manually determined cell outlines from the transmitted light image. To accurately quantify differences between samples and timepoints, cells were segmented from every image into individual ROIs using transmitted light images. Integrated NIR fluorescence intensity was calculated for each ROI and then divided by the cell area of the respective ROI. This resulted in normalized intensity measurements which were independent of cell size and number.

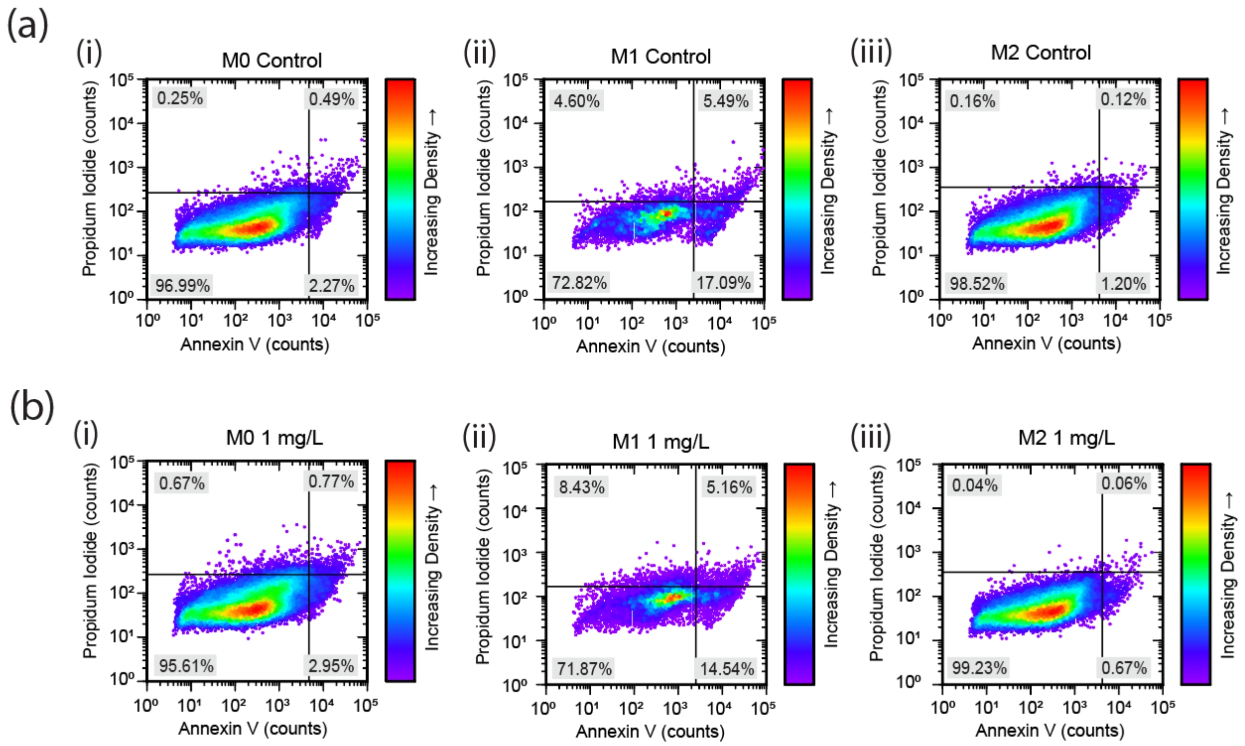


**Figure S4.** Cell viability assay to assess cytotoxic effects. (a) Scatter plots for Annexin V/PI viability assay with no SWCNTs. (b) Scatter plots for Annexin V/PI viability assay after 24 hours of continuous culture in a 1mg $L^{-1}$ $GT_6$-SWCNT solution.

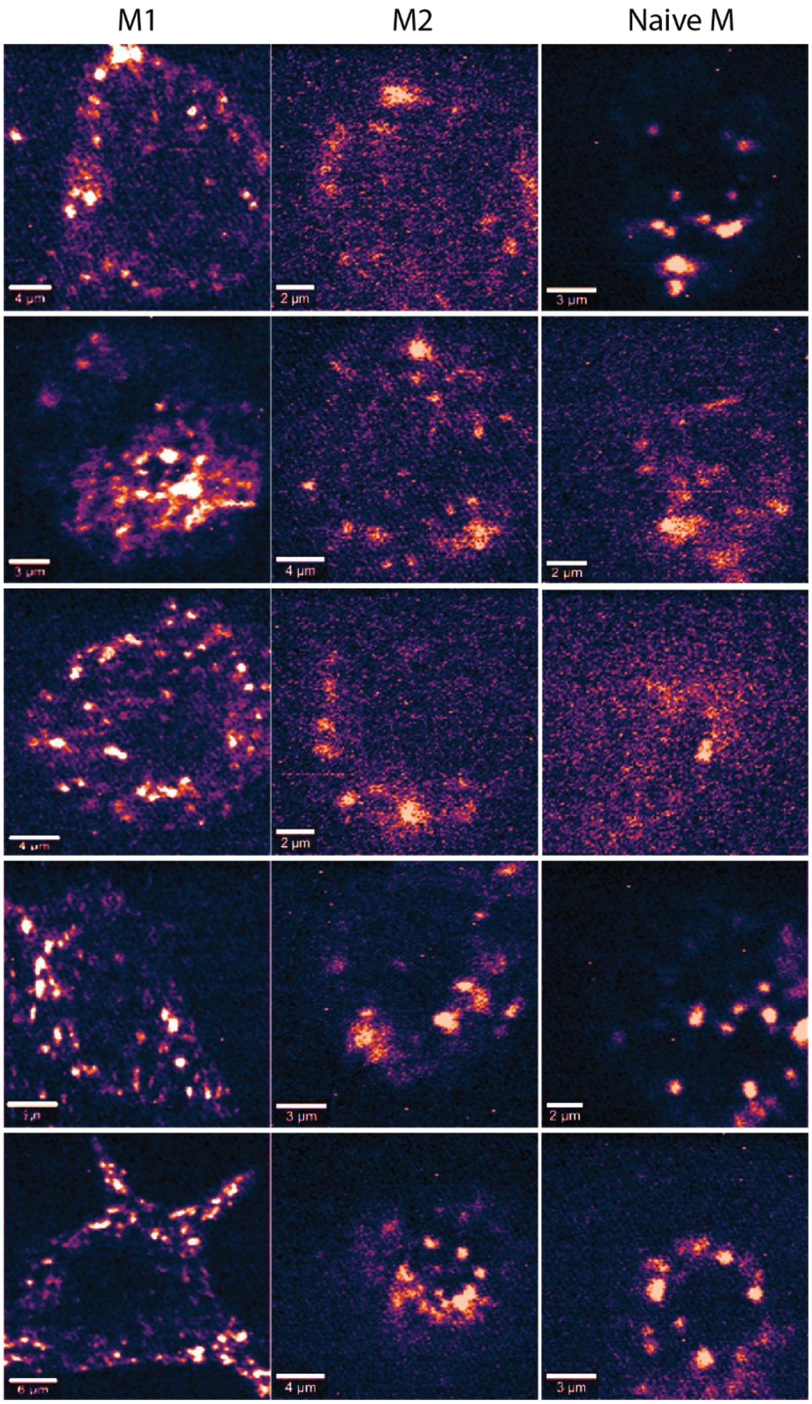


**Figure S5.** Additional confocal Raman microscope images. Integrated G-band Raman intensity images at 100x of all macrophage phenotypes, M1, M2 and naïve macrophages, imaged at 0.5 hours after SWCNT incubation.

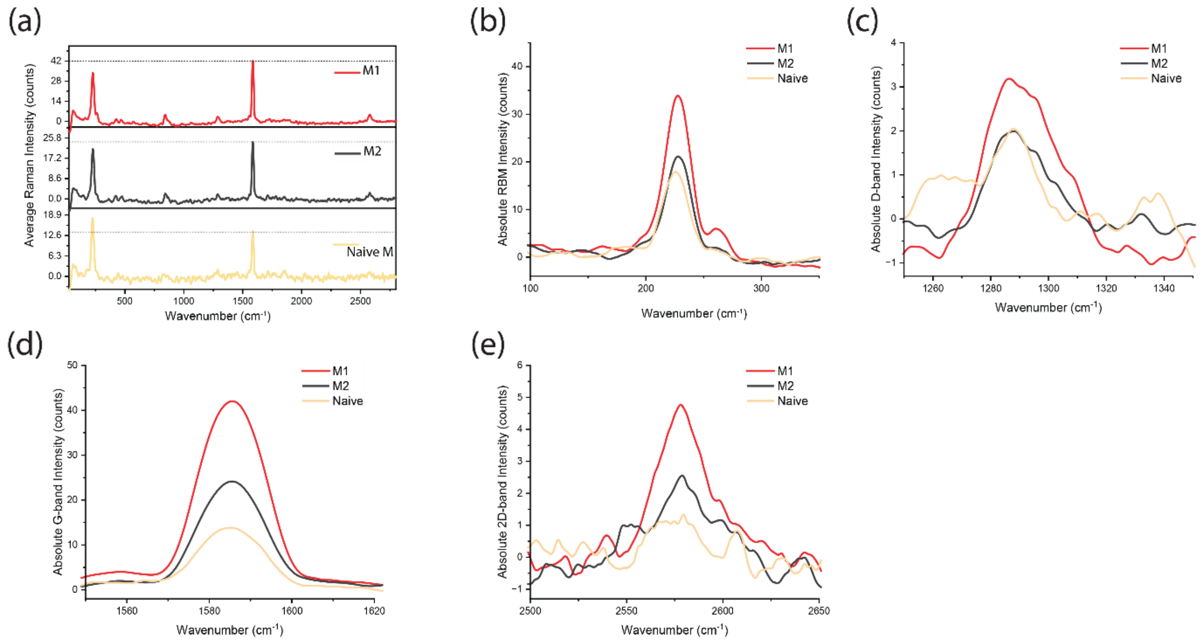


**Figure S6.** Confocal Raman microscopy analysis of SWCNT uptake by different macrophage phenotypes. (a) Stacked line plot showing full spectrum average Raman intensity of M1, M2 and naïve macrophages. Line plots showing differences in macrophage phenotypes among different SWCNT Raman features (b) Radial breathing mode (RBM), (c) D-band, (d) G-band, and (d) 2D-band.

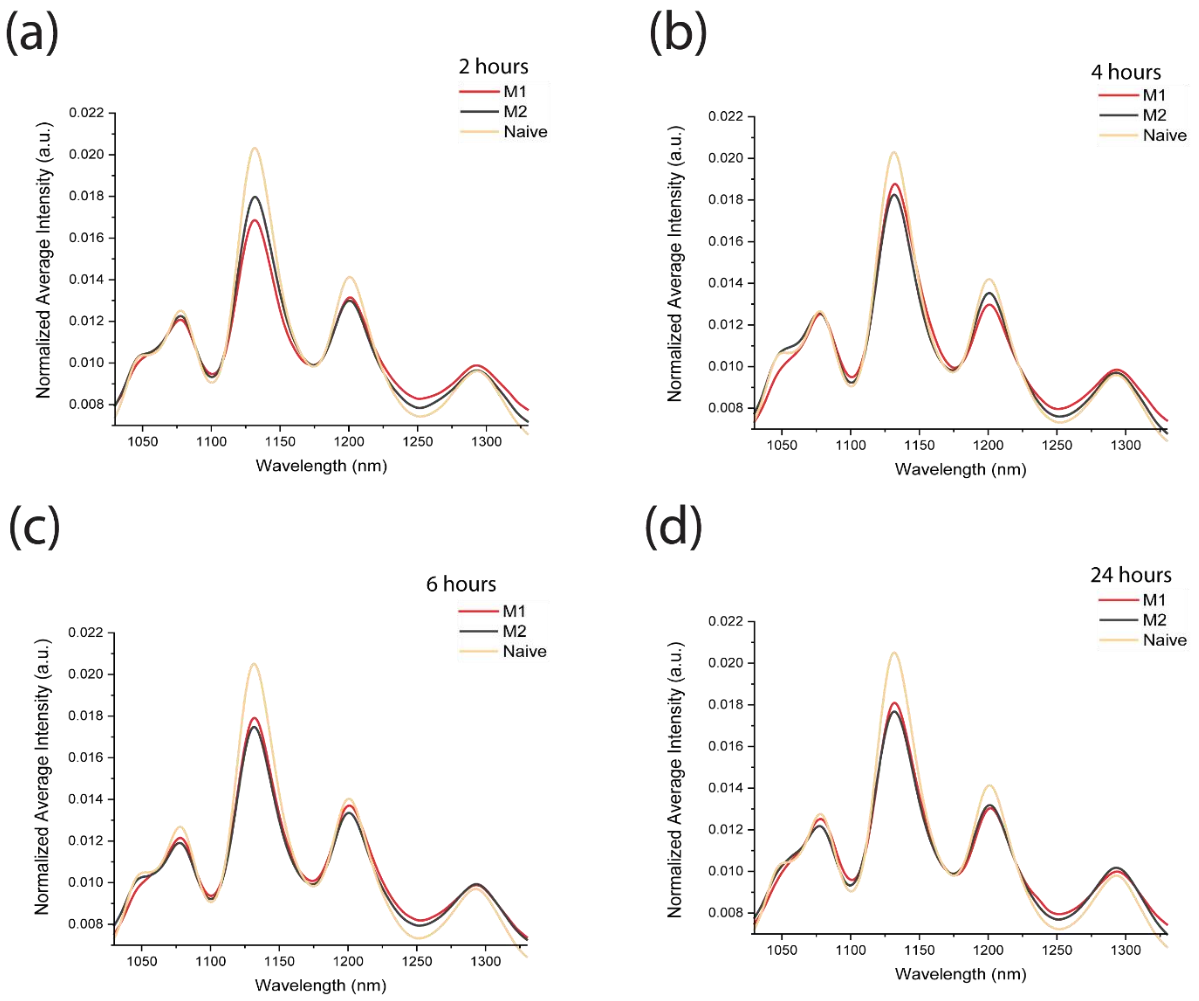


**Figure S7.** $GT_{15}$-SWCNT fluorescence response to different macrophage phenotypes, M1, M2, naïve macrophages across different time points (a) 2 hours (b) 4 hours (c) 6 hours (d) 24 hours. Y-axes represent the NIR fluorescence spectra averaged over all cells of that condition, normalized by the integrated intensity.

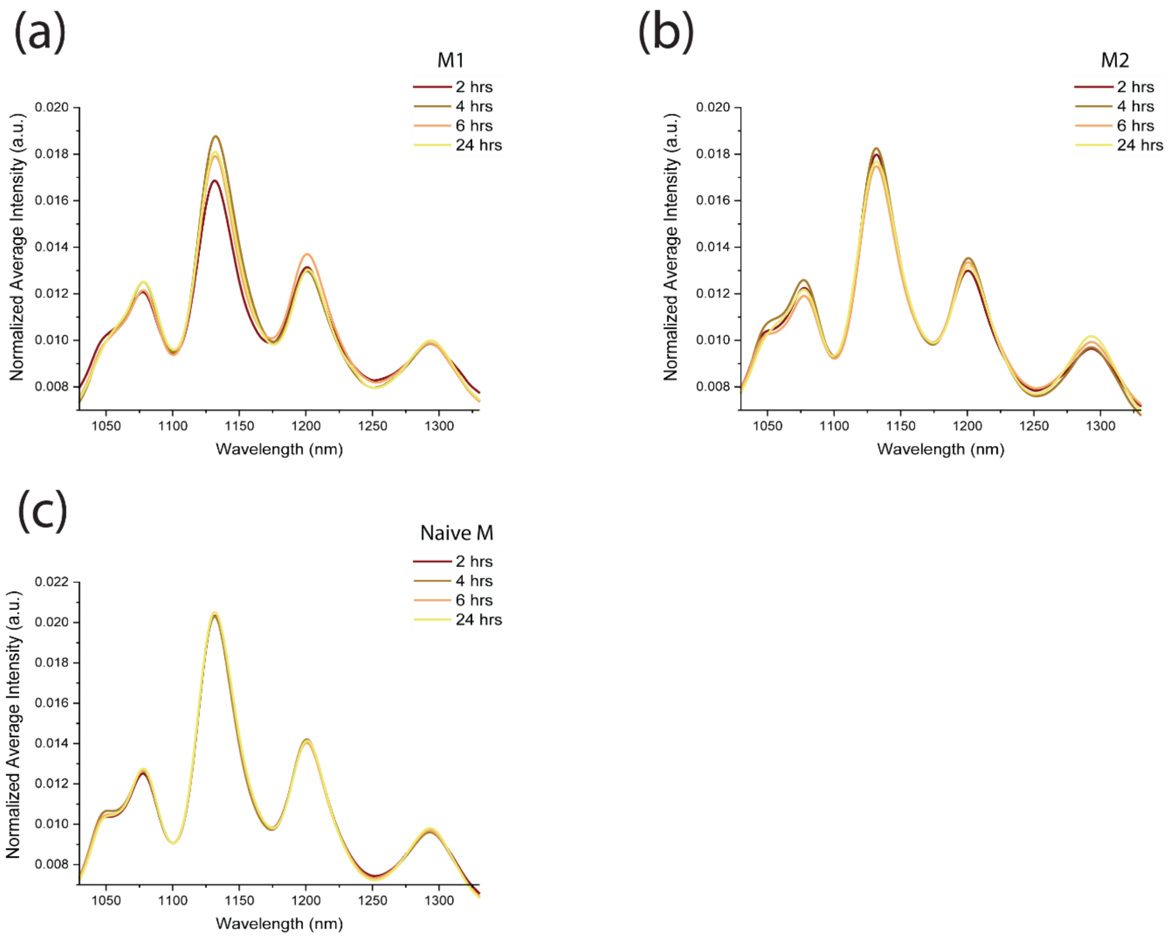


**Figure S8.** $GT_{15}$-SWCNT time dependent response comparison of each macrophage phenotype (a) M1, (b) M2, and (c) naïve macrophages across three timepoints of 2 hours, 4 hours and 6 hours. Y-axes represent the NIR fluorescence spectra averaged over all cells of that condition, normalized by the integrated intensity.

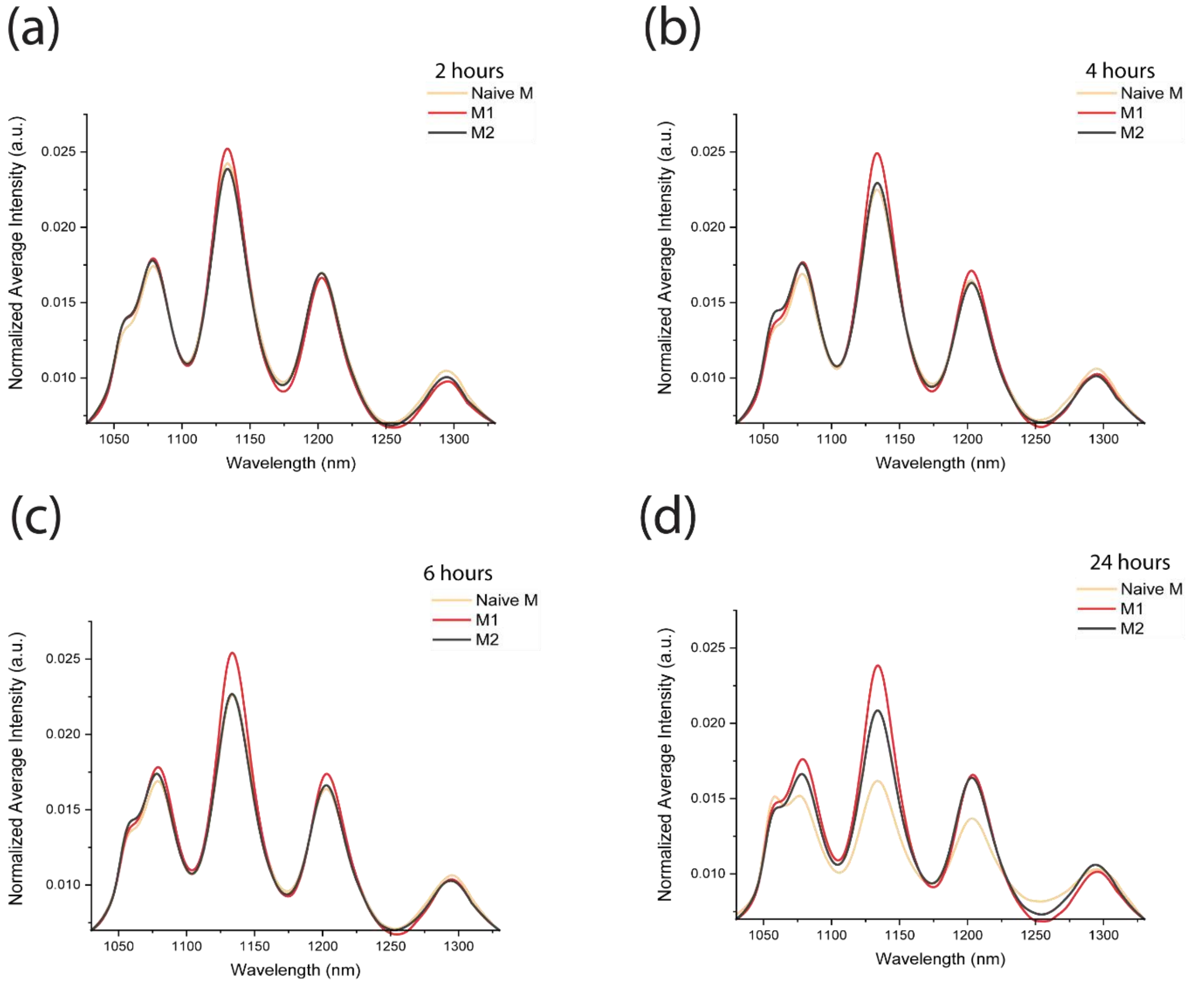


**Figure S9**. $GT_{30}$-SWCNT fluorescence response to different macrophage phenotypes, M1, M2, naïve macrophages across different time points (a) 2 hours (b) 4 hours (c) 6 hours (d) 24 hours. Y-axes represent the NIR fluorescence spectra averaged over all cells of that condition, normalized by the integrated intensity.

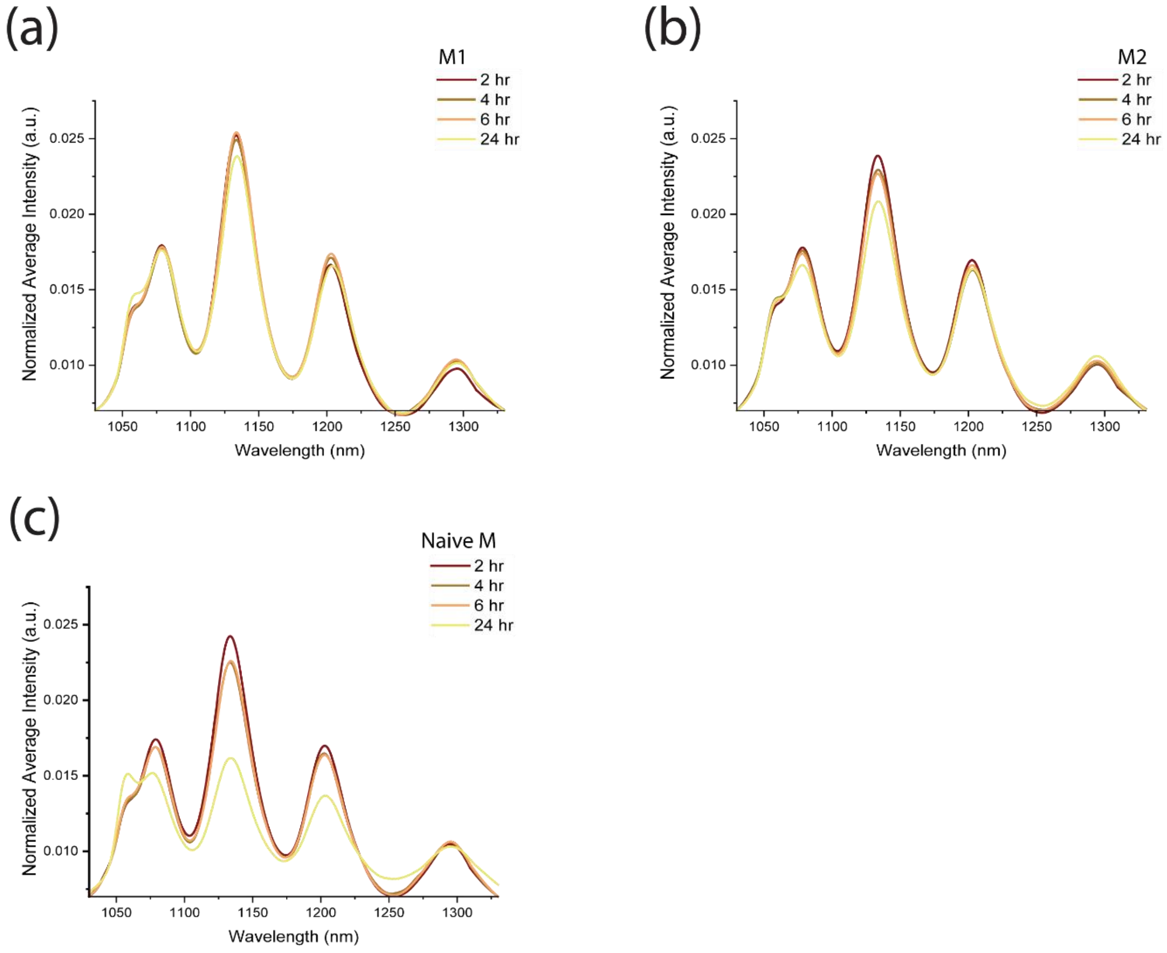


**Figure S10.** $GT_{30}$-SWCNT time dependent response comparison of each macrophage phenotype (a) M1 (b) M2 (c) naïve macrophages across three timepoints of 2 hours, 4 hours and 6 hours. Y-axes represent the NIR fluorescence spectra averaged over all cells of that condition, normalized by the integrated intensity.

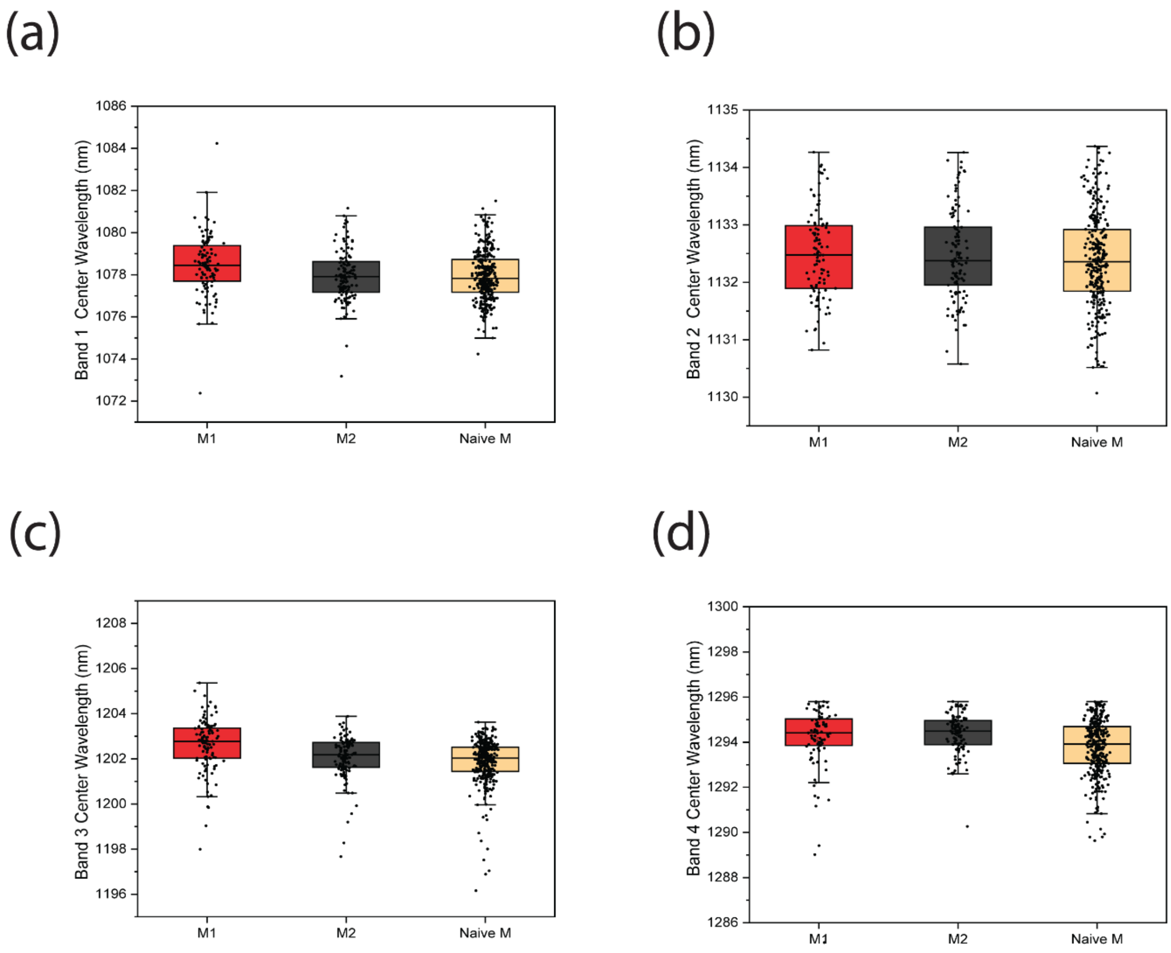


**Figure S11.** $GT_{15}$-SWCNT center wavelength comparison among different macrophage phenotypes, M1, M2, naïve macrophages across band containing different chiral species (a) band 1, (b) band 2, (c) band 3, and (d) band 4. Each ROI in the box and whisker plot is a single cell. Minimum of $n \geq 300$ cells per condition were used. Boxes represent 25–75% of the data, horizontal lines represent medians, and whiskers represent mean ± s.d.

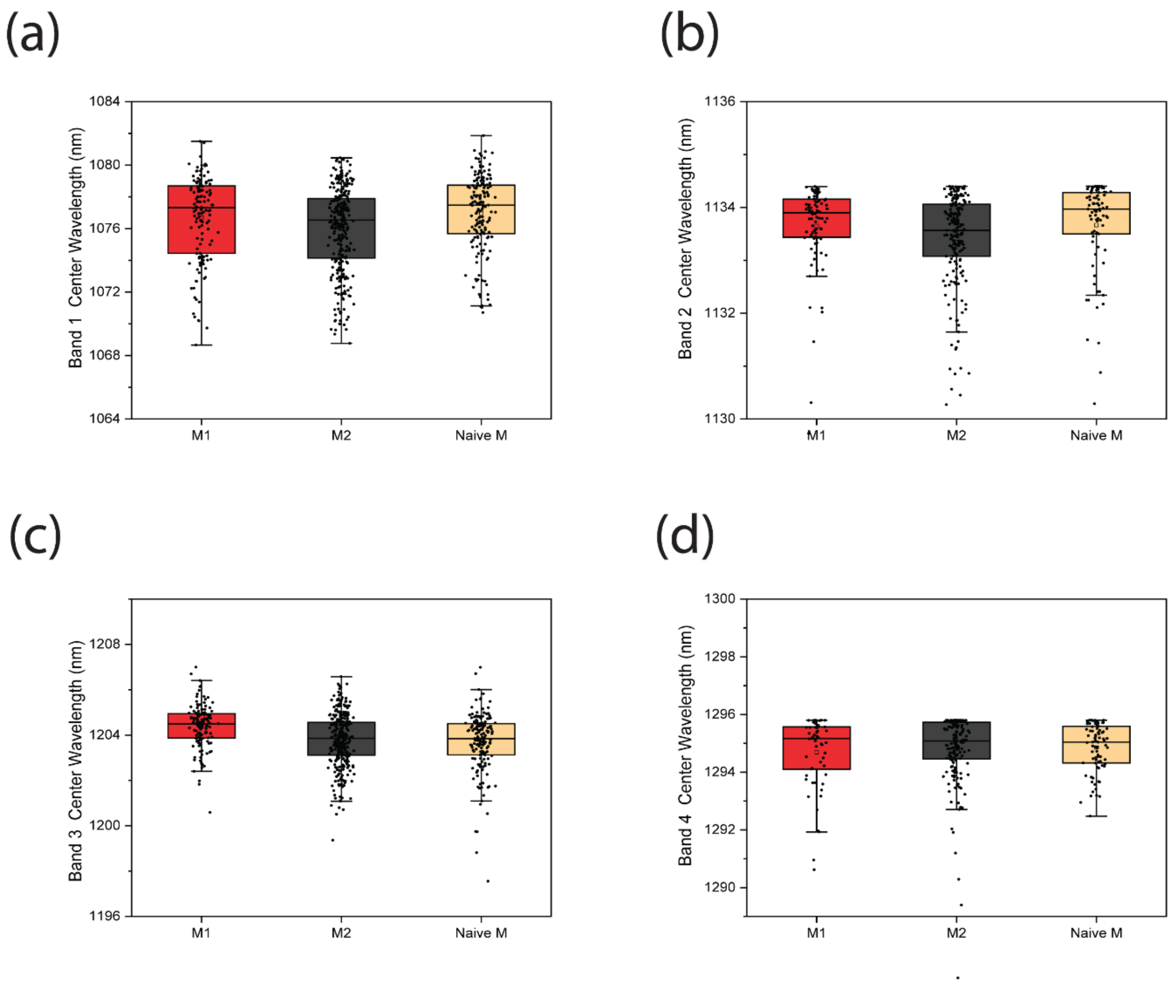


**Figure S12.** $GT_{30}$-SWCNT center wavelength comparison among different macrophage phenotypes, M1, M2, naïve macrophages across band containing different chiral species (a) band 1, (b) band 2, (c) band 3, and (d) band 4. Each ROI in the box and whisker plot is a single cell. Minimum of $n \geq 300$ cells per condition were used. Boxes represent 25–75% of the data, horizontal lines represent medians, and whiskers represent mean ± s.d.

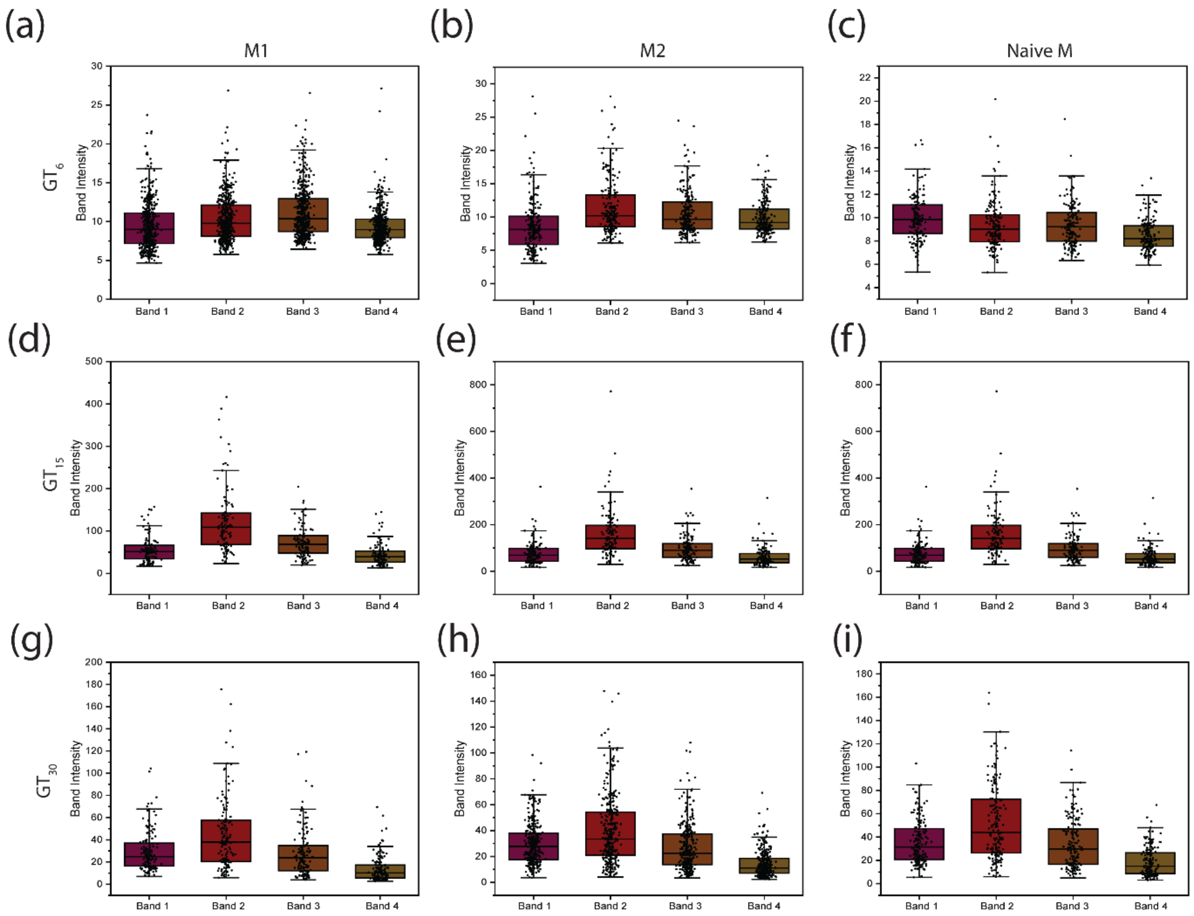


**Figure S13.** DNA-SWCNT band intensity response comparison. Box and whisker plots showing differences in band intensities among all macrophage phenotypes across all three DNA sequences (a) M1- $GT_6$, (b) M2- $GT_6$, (c) NM-$GT_6$, (d) M1-$GT_{15}$, (e) M2- $GT_{15}$, (f) NM-$GT_{15}$, (g) M1- $GT_{30}$, (h) M2- $GT_{30}$, and (i) NM-$GT_{30}$. Each ROI in the box and whisker plot is a single cell. Minimum of $n \geq 300$ cells per condition were used. Boxes represent 25–75% of the data, horizontal lines represent medians, and whiskers represent mean ± s.d.

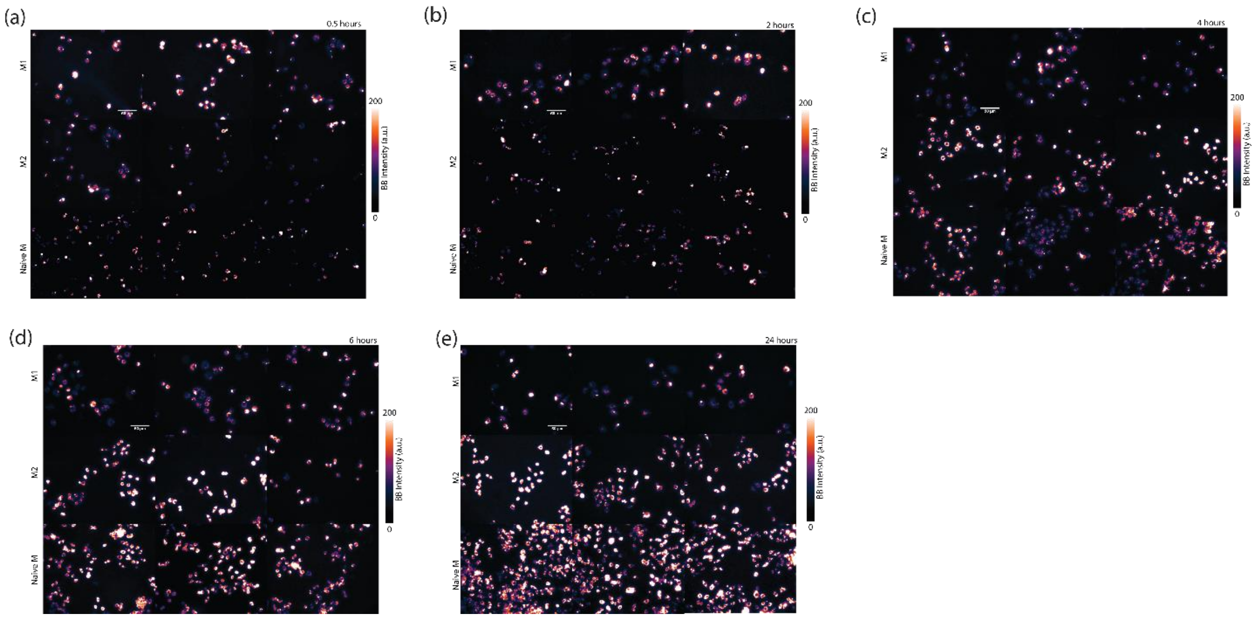


**Figure S14.** $GT_{15}$-SWCNT NIR fluorescence images of all macrophages phenotypes, M1, M2, naïve macrophages across all time points acquired in triplicates (a) 0.5 hours, (b) 2 hours, (c) 4 hours, (d) 6 hours, and (e) 24 hours. Scale bar on all images is 50 µm.

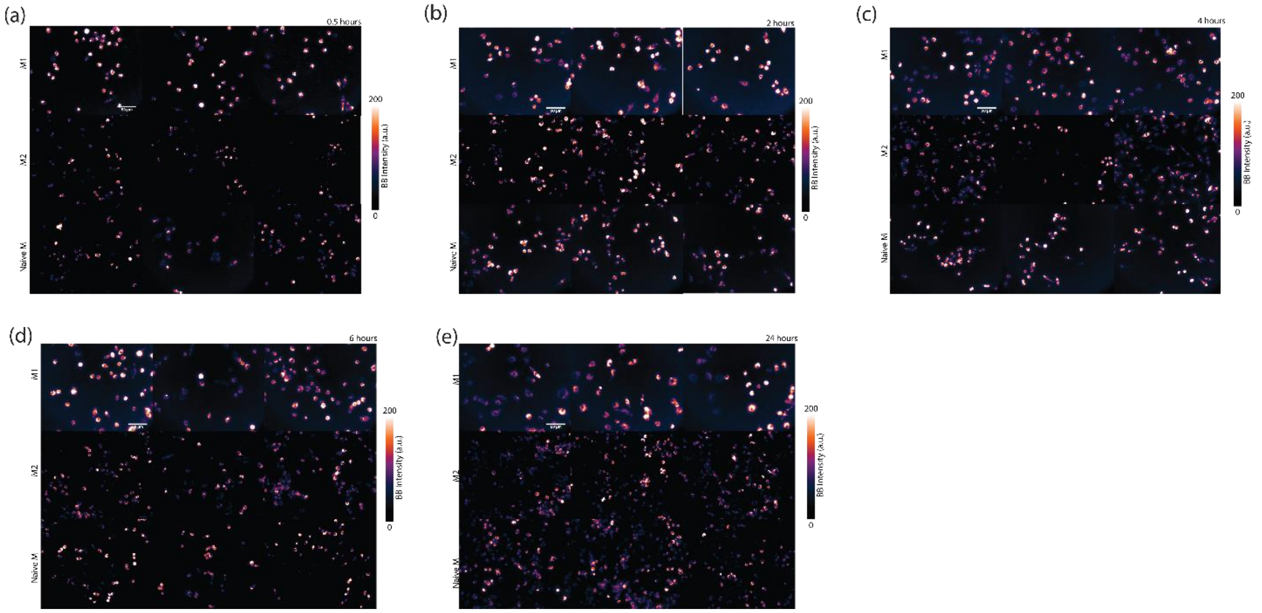


**Figure S15.** $GT_{30}$-SWCNT NIR fluorescence images of all macrophages phenotypes, M1, M2, naïve macrophages across all time points acquired in triplicates (a) 0.5 hours, (b) 2 hours, (c) 4 hours, (d) 6 hours, and (e) 24 hours. Scale bar on all images is 50 µm.

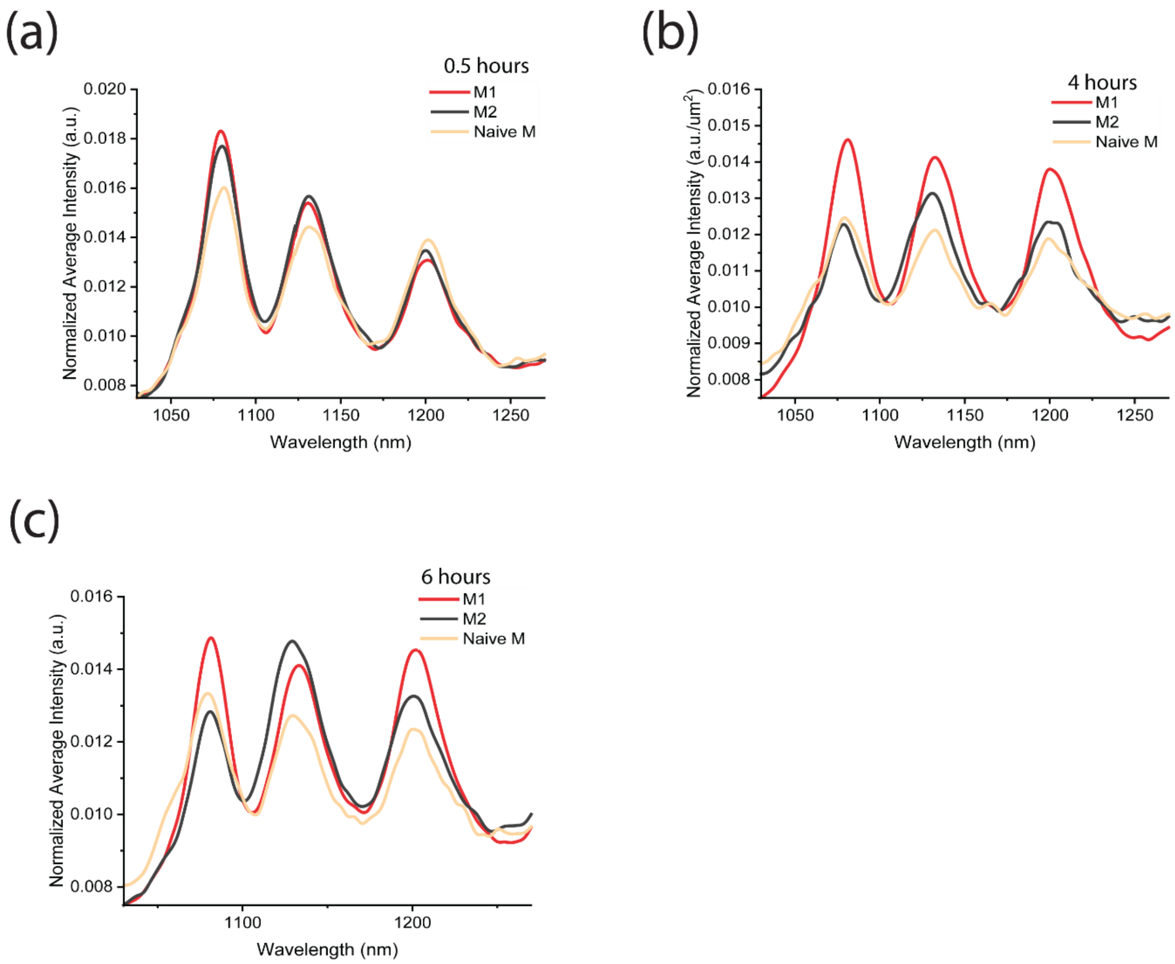


**Figure S16.** $GT_6$-SWCNT fluorescence response to different macrophage phenotypes, M1, M2, naïve macrophages across different time points (a) 0.5 hours (b) 4 hours (c) 6 hours. Y-axes represent the NIR fluorescence spectra averaged over all cells of that condition, normalized by the integrated intensity.

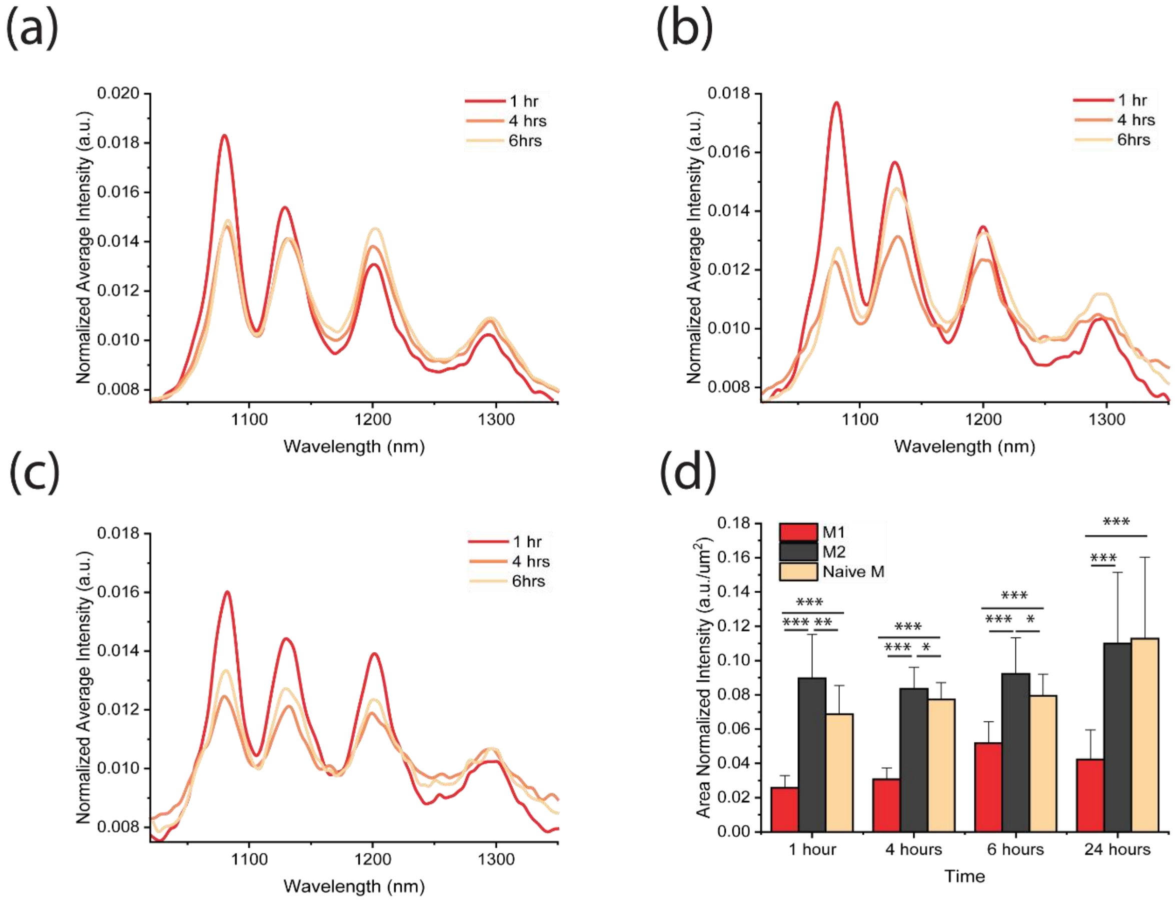


**Figure S17.** Time dependent response comparison of each macrophage phenotype (a) M1, (b) M2, and (c) naïve macrophages across three timepoints of 1 hour, 4 hours and 6 hours. (d) Bar graph showing the average area normalized intensity of all different macrophage phenotypes through all time points. Bars represent the average, and whiskers represent mean ± s.d. for each condition ($n \geq 300$ cells per condition).

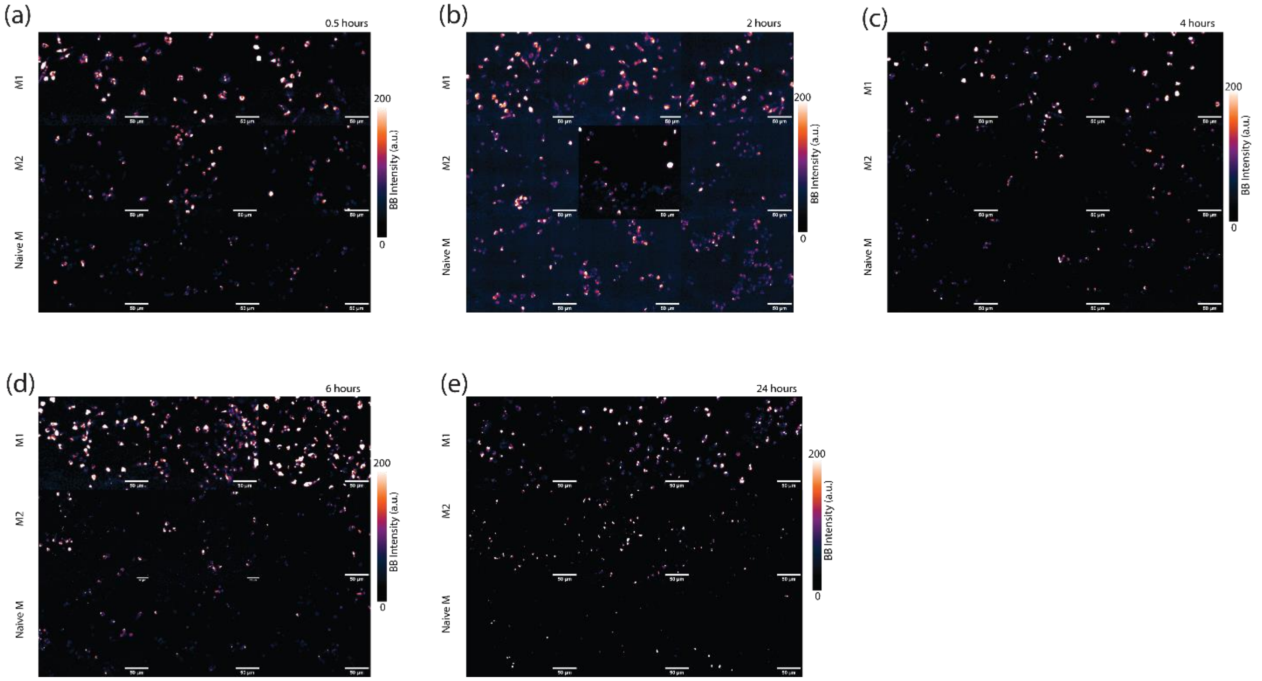


**Figure S18.** $GT_6$-SWCNT near-infrared fluorescence images of all macrophages phenotypes, M1, M2, naïve macrophages across all time points acquired in triplicates (a) 0.5 hours, (b) 2 hours, (c) 4 hours, (d) 6 hours, and (e) 24 hours. Scale bar on all images is 50 µm.

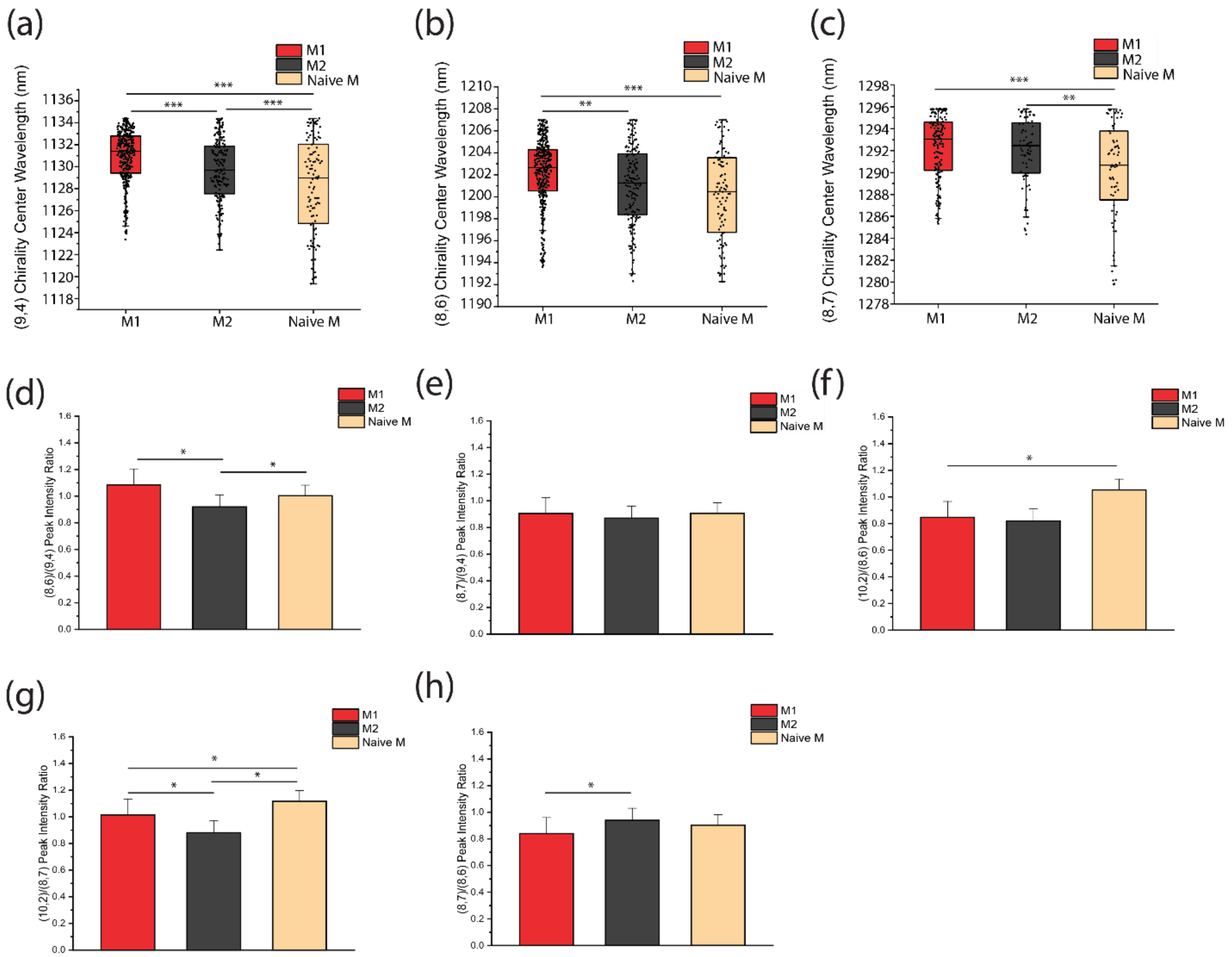


**Figure S19.** $GT_6$-SWCNT NIR fluorescence features used for machine learning. Box and whisker diagram for chiral species center wavelength (a) (9,4) (b) (8,6) (c) (8,7). Each ROI in the box and whisker plot is a single cell. Minimum of $n \geq 300$ cells per condition were used. Boxes represent 25–75% of the data, horizontal lines represent medians, and whiskers represent mean ± s.d. Two-Sample t-test hypothesis testing analysis was performed (***$p < 0.001$, **$p < 0.01$ and *$p < 0.05$). Bar graph showing peak intensity ratios (d) (8,6)/(9,4) (e) (8,7)/(9,4) (f) (10,2)/(8,6) (g) (10,2)/(8,7) (h) (8,7)/(8,6), for all cell phenotypes. Bars represent the average, and whiskers represent mean ± s.d. for each condition ($n \geq 300$ cells per condition). Two-Sample t-test hypothesis testing analysis was performed between different samples and between different time points. (*$p < 0.05$).

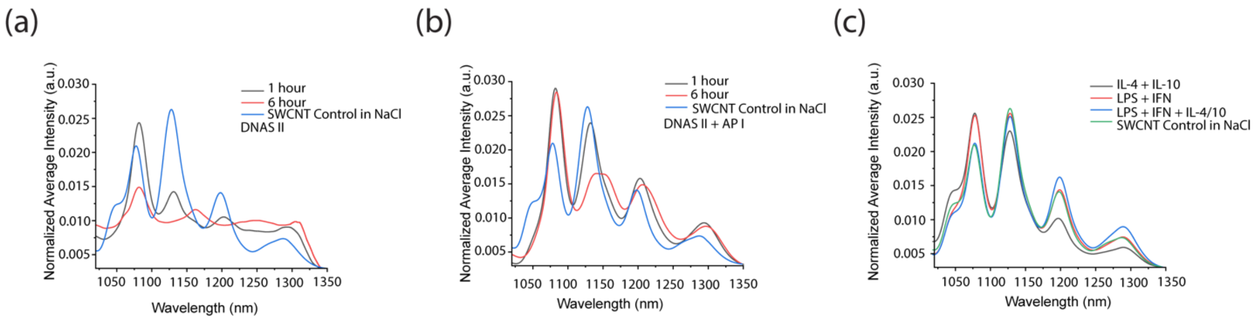


**Figure S20.** $GT_6$-SWCNT fluorescence response to (a) DNase II, (b) a mixture of DNase II and AP I at two biologically relevant time points and (c) mixture of M1 and M2 cytokines.

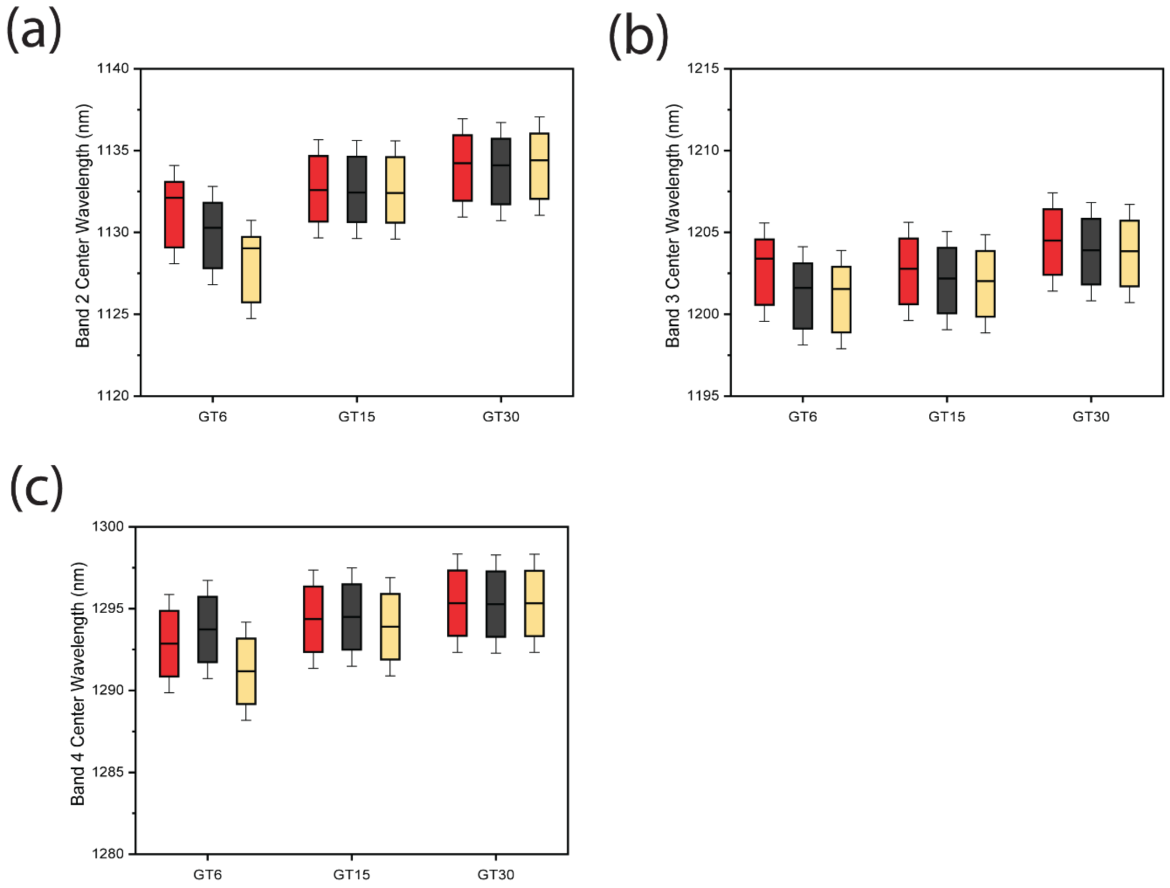


**Figure S21.** DNA-SWCNT center wavelength response comparison. Box and whisker plots showing differences in band center wavelength among all macrophage phenotypes M1, M2 and naïve macrophages across all three DNA sequences, $GT_6$, $GT_{15}$ and $GT_{30}$. (a) band 2, (b) band 3, and (c) band 4. Boxes represent 25–75% of the data, horizontal lines represent medians, and whiskers represent mean ± s.d.

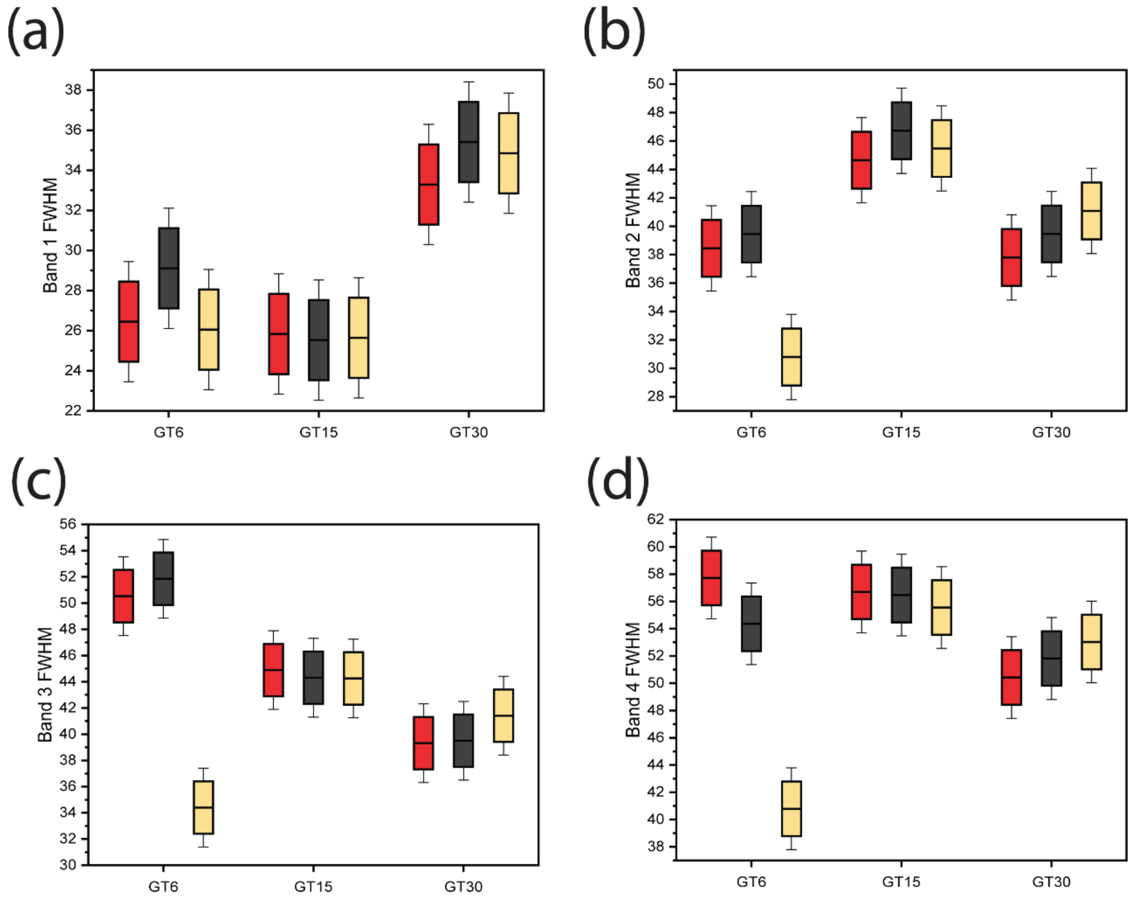


**Figure S22.** DNA-SWCNT full width half max response comparison. Box and whisker plots showing differences in full width at half maximum (FWHM) among all macrophage phenotypes M1, M2 and naïve macrophages across all three DNA sequences, $GT_6$, $GT_{15}$ and $GT_{30}$. (a) band 1, (b) band 2, (c) band 3, and (d) band 4. Boxes represent 25–75% of the data, horizontal lines represent medians, and whiskers represent mean ± s.d.

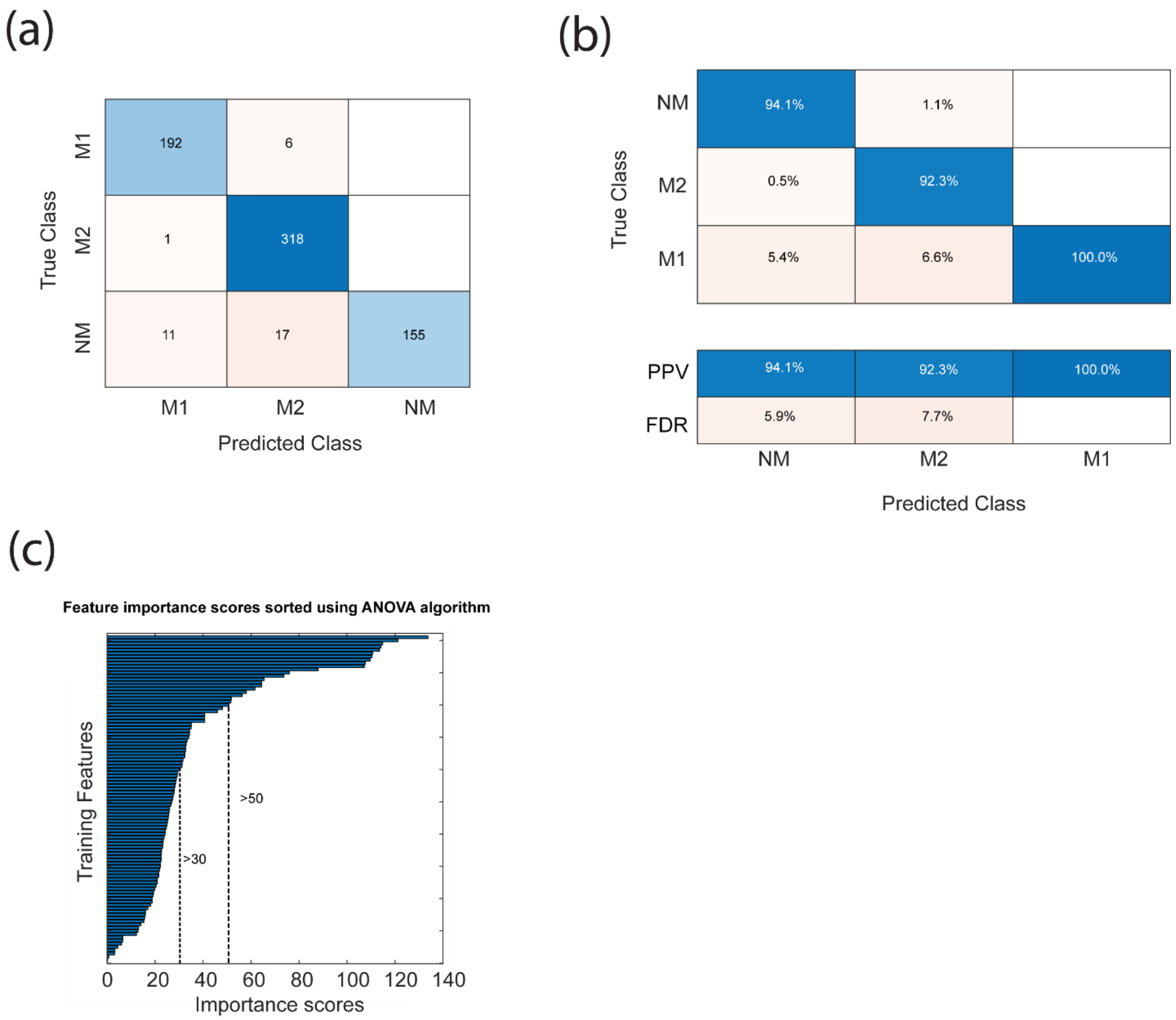


**Figure S23.** $GT_6$-SWCNT 6-hour Machine Learning data analysis (a) Confusion matrix of all macrophage types, M1, M2 and Naïve macrophages, for 6-hour input data. (b) Confusion matrix for all cell types showing percentages of true predictive values and false discovery rates. (c) Feature importance distribution graph built using an ANOVA algorithm for Machine Learning.

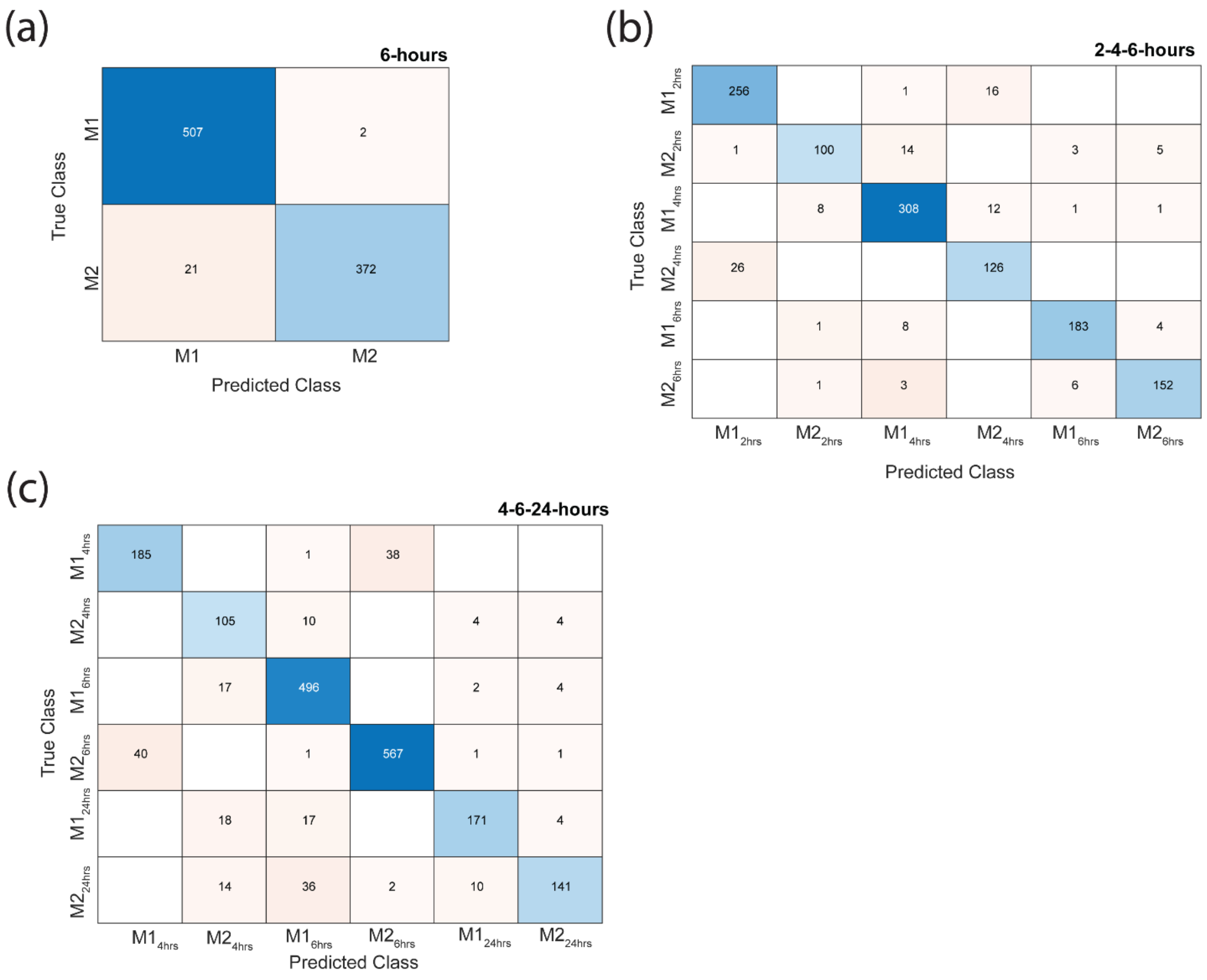


**Figure S24.** $GT_6$-SWCNT Machine Learning confusion matrices of M1 and M2 macrophages for different input data sets (a) 6 hours, (b) 2, 4, and 6 hours, (c) 4, 6, and 24 hours. The number of cells in each data set is > 100 cells per condition.

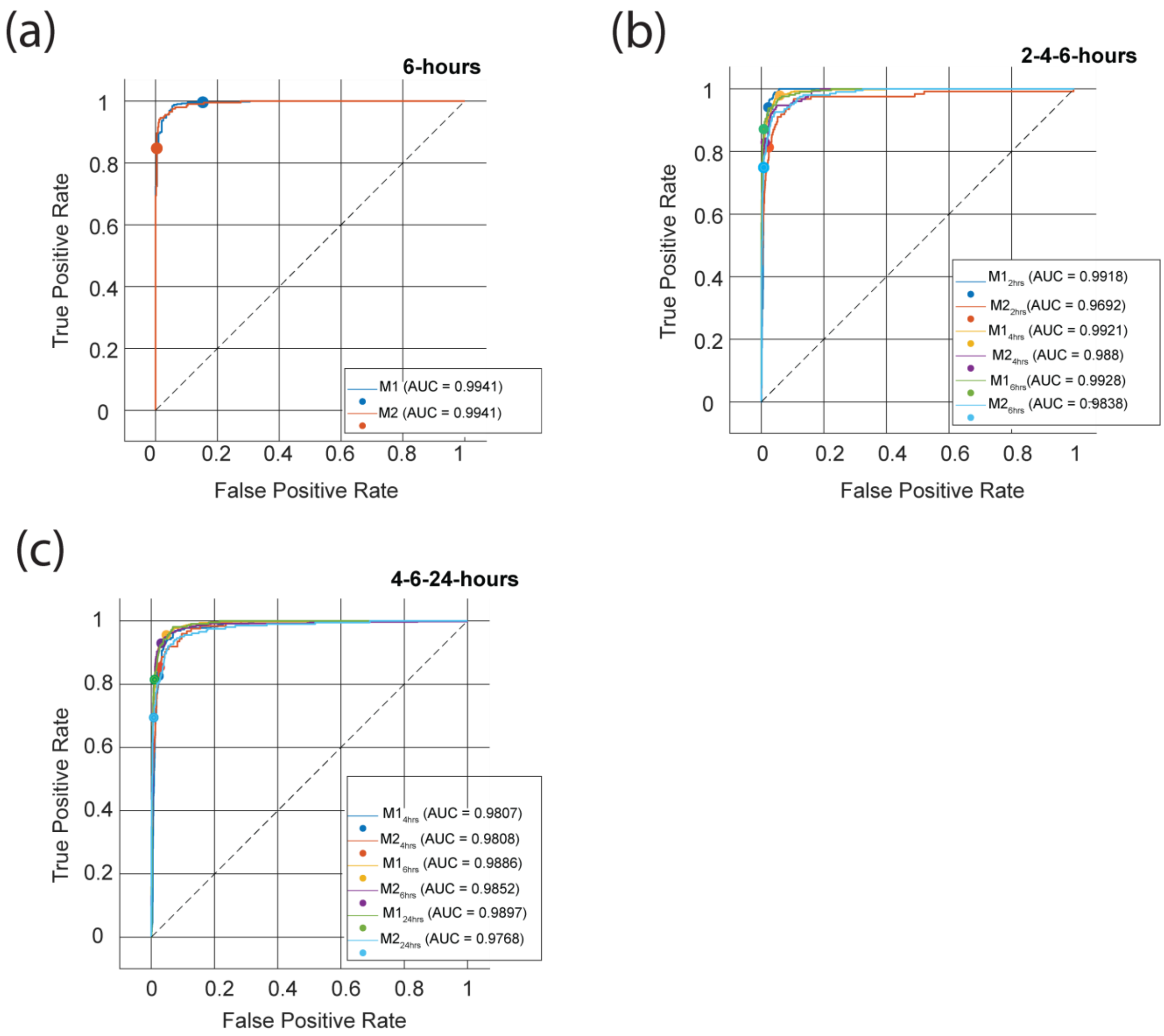


**Figure S25.** $GT_6$-SWCNT Machine Learning ROC curves of M1 and M2 macrophages for different input data sets (a) 6 hours, (b) 2, 4, and 6 hours, and (c) 4, 6, and 24 hours.

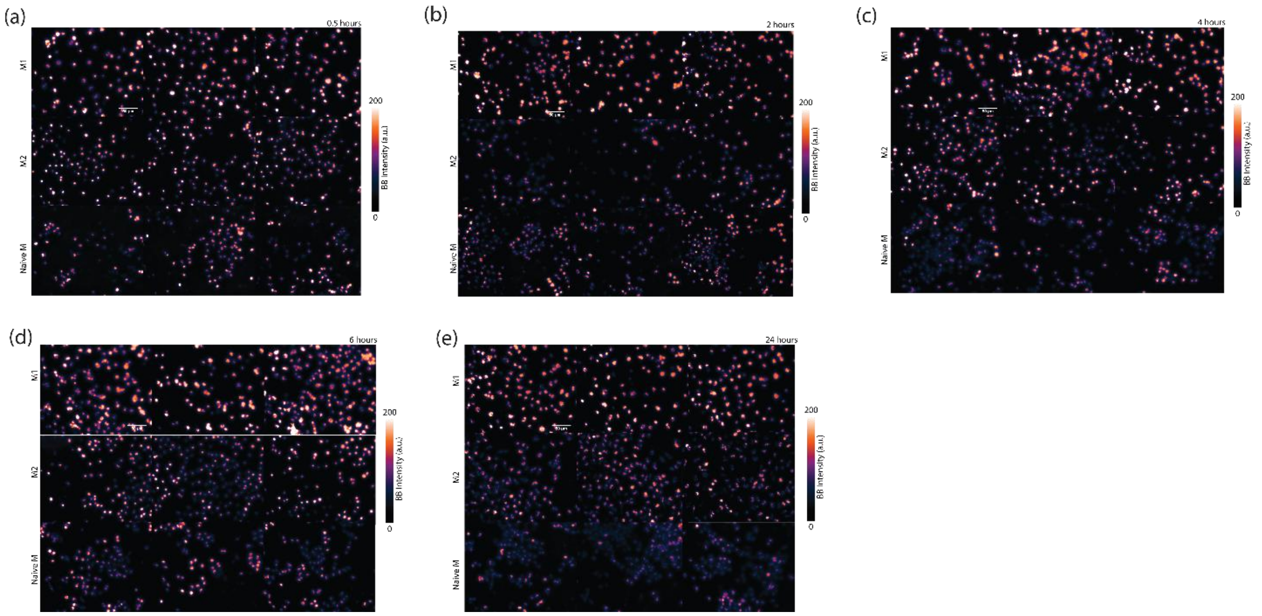


**Figure S26.** $GT_6$-SWCNT NIR fluorescence images of all primary macrophage phenotypes, M1, M2, naïve bone marrow derived macrophages across all time points acquired in triplicates (a) 0.5 hours, (b) 2 hours, (c) 4 hours, (d) 6 hours, and (e) 24 hours. Scale bar on all images is 50 µm.

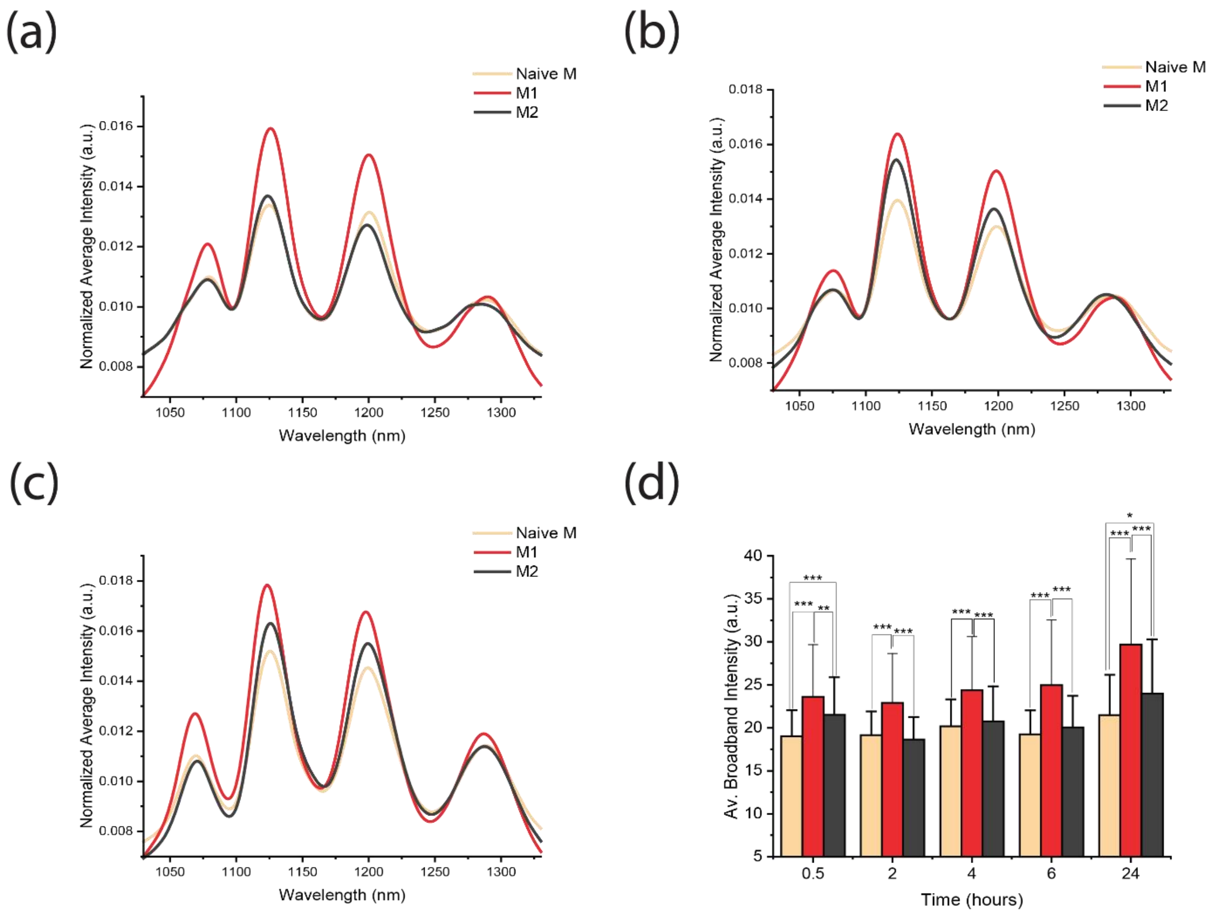


**Figure S27.** $GT_6$-SWCNT fluorescence response to different primary bone marrow derived macrophages (BMDM) phenotypes, M1, M2, naïve BMDM across different time points (a) 4 hours, (b) 6 hours, and (c) 24 hours. (d) Bar graph showing the average area normalized intensity of all different macrophage phenotypes through all time points. Bars represent the average, and whiskers represent mean ± s.d. for each condition ($n \geq 300$ cells per condition). Two-Sample t-test hypothesis testing analysis was performed between different samples and between different time points. (*$p < 0.05$).

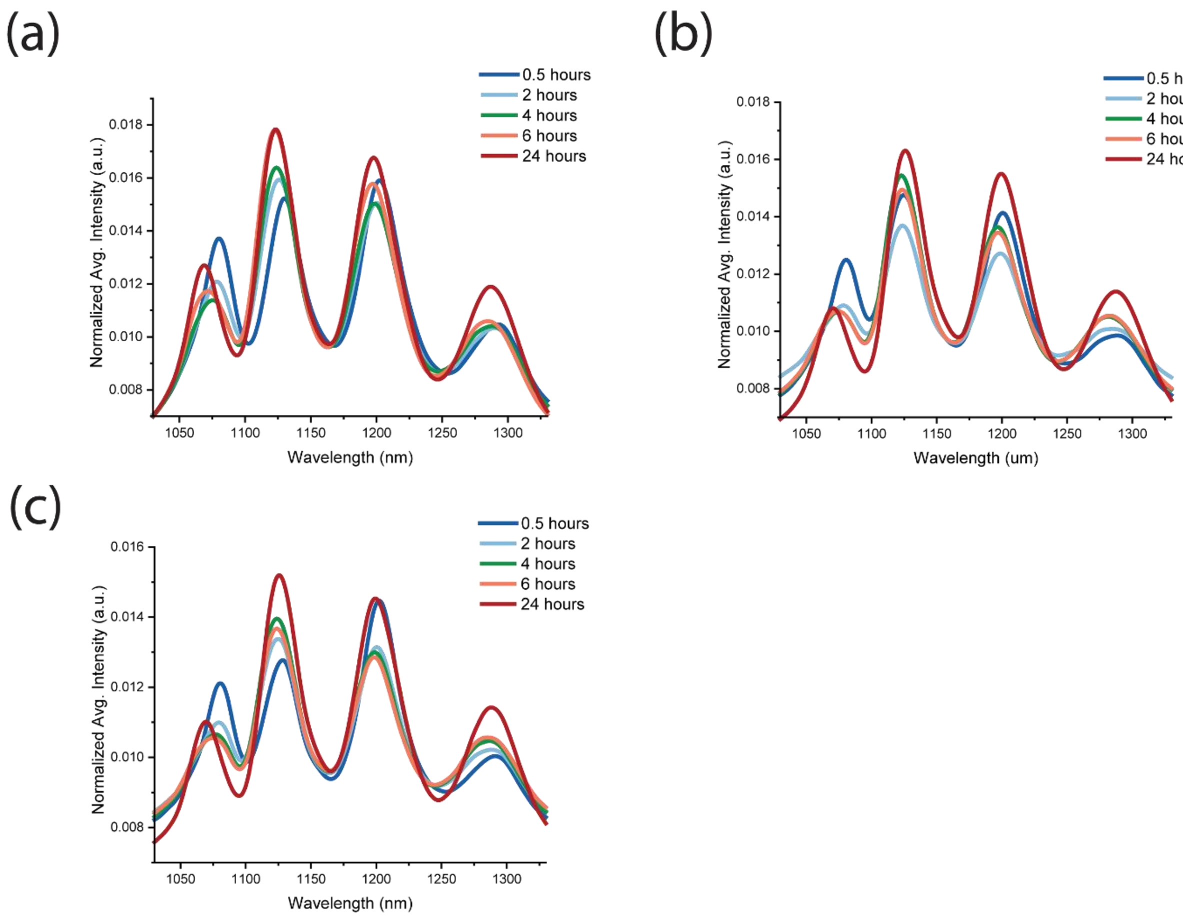


**Figure S28.** $GT_6$-SWCNT time dependent response comparison of each primary macrophage phenotypes (a) M1 (b) M2 (c) naïve BMDM across five timepoints of 0.5 hour, 2 hours, 4 hours, 6 hours and 24 hours.
